# ER-associated degradation relying on protein O-mannosylation

**DOI:** 10.1101/2025.01.15.633243

**Authors:** Leticia Lemus, Hadar Meyer, Ana Rodriguez, Maya Schuldiner, Veit Goder

**Affiliations:** Dept. of Genetics, University of Seville, Av. Reina Mercedes 6, 41012 Seville, Spain; Dept. of Molecular Genetics, Meyer Bldg. Room 122, Weizmann Institute of Sciences, 76100 Rehovot, Israel

**Keywords:** Protein O-mannosylation, ERAD, ER quality control, Pbn1, Gpi14

## Abstract

Protein quality control in the secretory pathway, whose dysfunction is linked to various human diseases, begins in the endoplasmic reticulum (ER) and involves ER-associated degradation (ERAD) of terminally misfolded species. We have addressed the roles of protein O-mannosylation in ERAD using complementary genome-wide screens in yeast. Our findings reveal that protein O-mannosylation by the conserved ER-resident O-mannosyltransferase complex Pbn1-Gpi14 generates ERAD cues. This mechanism participates in regulating ERAD of substrates that contain serine-rich regions and provides a fail-safe mechanism for the degradation of non-N-glycosylated misfolded proteins. Our data suggest that the *de novo* synthesis of ERAD cues through protein O-mannosylation is an essential component of ER quality control.

## Introduction

Folding of proteins in the secretory pathway starts after their translocation across or integration into the membrane of the endoplasmic reticulum (ER) and is monitored by ER protein quality control (ERQC), a process that is essential for cellular homeostasis (Ellgaard and Helenius, 2003). ERQC directs non-aggregated terminally misfolded polypeptides to ER-associated protein degradation (ERAD), which includes their retrotranslocation into the cytosol by ER membrane-embedded protein complexes followed by their ubiquitination and delivery to the proteasome (Hiller et al., 1996; Werner et al., 1996; Swanson et al., 2001; Denic et al., 2006; Carvalho et al., 2006; Nakatsukasa et al., 2008; Biederer et al., 1997; Jarosch et al., 2002; Stein et al., 2014; Bodnar and Rapoport, 2017; Kim et al., 2004).

Most proteins in the secretory pathway undergo N-glycosylation within the ER, playing crucial roles in ERQC and ERAD. Initially, preassembled polyglycan structures are added to site-specific asparagines in luminal protein domains (Helenius and Aebi, 2004; Ruiz-Canada et al., 2009). Subsequent enzymatic glycan trimming is coupled to protein folding, generating N-glycan structures that either promote (re-)binding to lectin-type chaperones for folding or target them for delivery to ERAD (Hammond et al., 1994; Caramelo et al., 2004; Molinari et al., 2004; Quan et al., 2008; Clerc et al., 2009). Although misfolded proteins lacking N-glycans can also be efficiently routed to ERAD (Leto et al., 2019), it remains unknown if and how these substrates are selectively labeled by other mechanisms.

Protein O-glycosylation, also known as O-mannosylation, is a second type of glycosylation that occurs within the ER. Single mannoses are attached primarily to serines or threonines by evolutionarily conserved, ER-resident protein mannosyltransferases (PMTs) (Haselbeck and Tanner, 1983; Lommel and Strahl, 2009; Neubert et al., 2016), as well as by distantly related transmembrane and tetratricopeptide repeat-containing proteins 1-4 (TMTC 1-4) (Larsen et al., 2017). PMTs are part of the GT-C superfamily of ER-resident mannosyltransferases, which also includes enzymes responsible for transferring mannose units for the synthesis of N-glycan precursors and glycosylphosphatidylinositol (GPI) anchors. All ER-resident mannosyltransferases use dolichol phosphate-mannose as a donor and share common catalytic motifs, as well as a common multi-transmembrane domain (TMD) architecture (Liu and Mushegian, 2003; Albuquerque-Wendt et al., 2019; Bloch et al., 2020; Alexander and Locher, 2023). PMTs have been linked to ERQC and are thought to possess protein chaperone functions through their MIR (protein O-mannosyltransferase [PMT], inositol 1,4,5-trisphosphate receptor [IP3R], and ryanodine receptor [RyR]) domains (Fukuda et al., 2001; Bai et al., 2019; Chiapparino et al., 2020). However, the role of protein O-mannosylation in ERQC remains unclear. Drug-induced inhibition of protein N-glycosylation has been found to increase global protein O-mannosylation and reduce protein aggregation (Harty et al., 2001; Nakatsukasa et al., 2004). In non-stressed yeast cells, Pmt1 and Pmt2 promote ER retention of misfolded proteins and physically interact with central ERAD components (Goder and Melero, 2011). Finally, protein O-mannosylation has been shown to block (re)binding to ER chaperones in vitro and is proposed to be a mechanism to terminate energy-consuming folding cycles in the ER in vivo (Xu et al., 2013).

The misfolded yeast protein Gas1* has been used as a model to study ERAD of GPI-anchored proteins (GPI-APs) (Fujita et al., 2006). Gas1* is a mutant form of Gas1, an abundant protein in the secretory pathway. Unlike Gas1, which is moderately O-mannosylated by Pmt4, Gas1* is highly O-mannosylated, primarily by the Pmt1/2 complex, making it an ideal model to study the roles of protein O-mannosylation in relation to ERQC and ERAD (Hirayama et al., 2008; Goder and Melero, 2011; Sikorska et al., 2016). The degradation of Gas1* is complex and occurs through distinct cellular pathways. A fraction is retained within the ER and subsequently routed to ERAD, while a larger pool is rapidly exported from the ER and targeted to the vacuole for degradation (Fujita et al., 2006; Hirayama et al., 2008; Goder and Melero, 2011; Sikorska et al., 2016; Lemus et al., 2021). Protein O-mannosylation contributes to ER retention of Gas1* and increases the fraction routed to ERAD, but its significance in this process remained unclear (Hirayama et al., 2008; Goder and Melero, 2011; Sikorska et al., 2016).

We performed complementary genome-wide screens in the yeast model *S. cerevisiae* to systematically address the roles of protein O-mannosylation in Gas1* degradation. Our results revealed that the evolutionarily conserved ER-resident proteins Pbn1 and associated Gpi14 generate ERAD cues through protein O-mannosylation. Previously, Pbn1-Gpi14 was known for forming GPI-mannosyltransferase I (GPI-MT I), responsible for transferring the first mannose onto the glycolipid glucosamine-(acyl)phosphatidylinositol (GlcN-(acyl)PI) during GPI anchor precursor biosynthesis (Ashida et al., 2005). Our data indicate that the protein O-mannosylation activity of Pbn1-Gpi14 is crucial for regulating ERAD of proteins with serine-rich regions and provides a fail-safe mechanism for the degradation of non-N-glycosylated misfolded proteins.

## Results

### Genome-wide yeast screens identify Pbn1 as a regulator for the degradation of misfolded GPI-AP Gas1*

We employed complementary genome-wide yeast screens to identify factors involved in ERAD of Gas1*. For a growth-based screen, we generated the chimeric protein Leu2-TMD- Gas1, composed of the normally short-lived Gas1* fused to Leu2 via the transmembrane domain (TMD) of ER-resident Ted1 (Fig. 1A). In cells lacking genomic *LEU2* (Δleu2), growth in the absence of leucine is predicted to occur only if the reporter fails to be degraded, such as upon genetic inhibition of ERAD (Fig. 1B). Consistent with this prediction, the ERAD mutant *Δubc7*, but not wild-type cells, supported growth in the absence of leucine (suppl. Fig. S1A). Leu2-TMD-Gas1* was transformed into a collection of strains containing viable individual deletions and decreased abundance by mRNA perturbation (*DAmP*) alleles of essential genes, resulting in over 5800 transformants covering approximately 93% of all genes (Fig. 1B). Transformants were replica-plated into media containing or lacking leucine, leading to the identification of 213 genes that affected reporter degradation, 36 of which encoded annotated ER proteins (Figs. 1B and E, suppl. Fig. S1B, and Table S1). Among these were all components of the Hrd1 complex, an ERAD machinery known to be involved in Gas1* degradation, confirming the screen’s functionality (Fig. 1E) For a complementary screen, we utilized GFP-Gas1* (Fig. 1C and Sikorska et al., 2016). The goal of this screen was to identify cellular factors whose overexpression leads to elevated degradation of GFP-Gas1, thereby reducing cellular GFP levels. GFP-Gas1 was first integrated into *Δted1* cells to increase its routing to ERAD (suppl. Fig. S1C and Sikorska et al., 2016), and then crossed with a collection of strains expressing individual ORFs with N-terminally fused mCherry under the control of the constitutively strong *TEF2* promoter. This process yielded more than 5900 double fluorescent strains, covering approximately 95% of all genes (Fig. 1D). Automated image acquisition and fluorescence signal quantification identified 153 genes that increased reporter degradation upon strong expression, 10 of which were predicted or verified ER protein-encoding genes (Figs. 1D and E, and suppl. Table S1). The essential gene PBN1 was the only gene identified in both screens (Fig. 1E).

**Figure 1.**
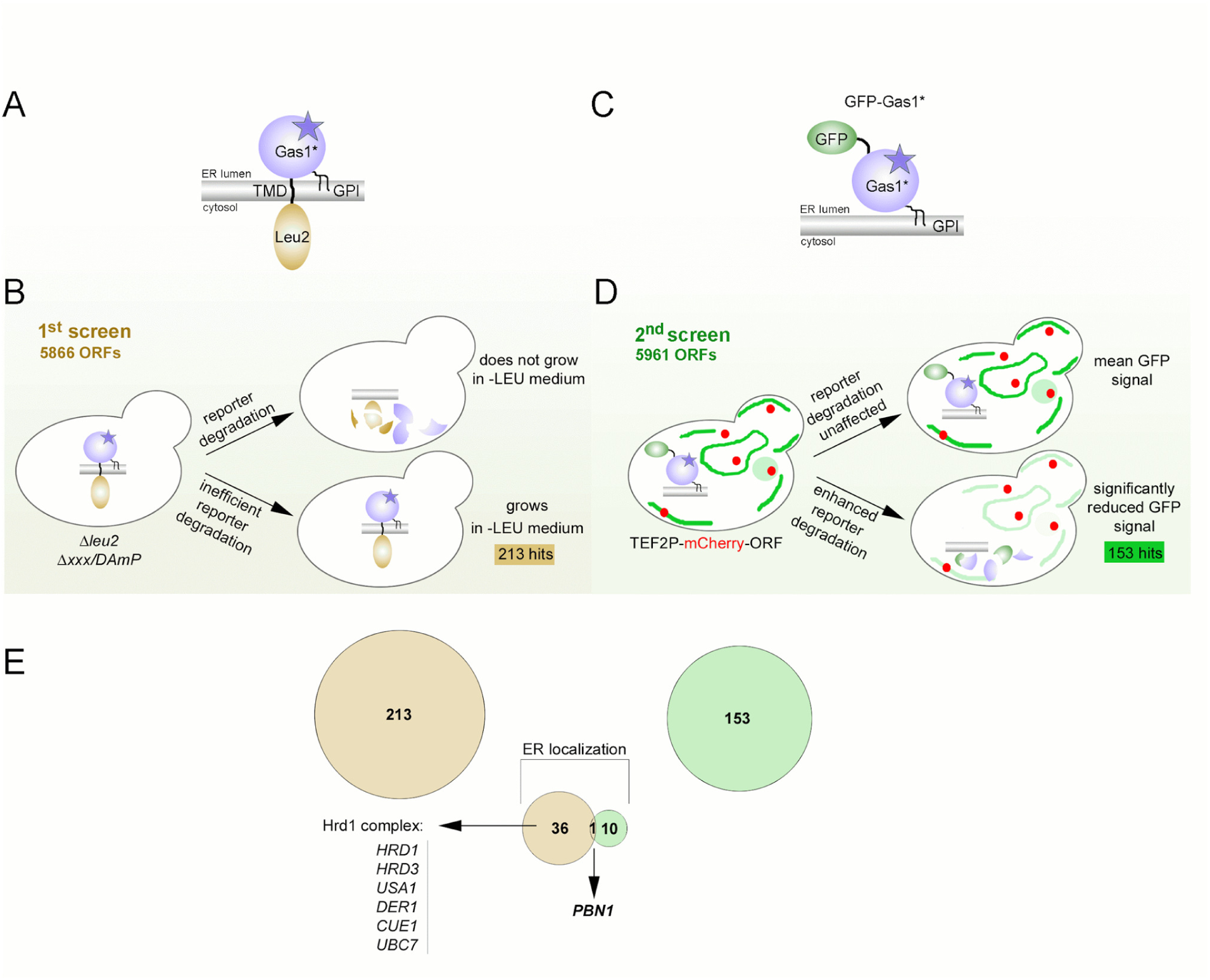
Genome-wide yeast screens identify Pbn1 as a regulator for the degradation of misfolded GPI-AP Gas1*. **(A)** Illustration of the Leu2-TMD-Gas1* reporter and its topology in the ER membrane. Yeast Leu2 was fused to the N-terminus of Gas1* via the transmembrane domain (TMD) of the ER-resident protein Ted1. Gas1* contains a C-terminal GPI anchor (GPI). (**B)** Schematic illustration of the genome-wide screen using Leu2-TMD- Gas1*. The collections of individual deletion mutants (*Δxxx*) and decreased abundance by mRNA perturbation (*DAmP*) mutants are auxotroph for leucine (*Δleu2*). They were transformed with a URA3-bearing plasmid expressing the reporter from the weak *GAL4* promoter. Transformants were grown in 96-well plates containing medium lacking uracil or both uracil and leucine. Efficient inhibition of Leu2-TMD-Gas1* degradation allowed cells to grow in medium lacking leucine (hits). (**C)** Illustration of the GFP-Gas1* reporter and its topology in the ER membrane. (**D)** A yeast query strain expressing genome-integrated GFP-Gas1* from its endogenous promoter was crossed with the collection of mutants expressing N-terminal mCherry fusions to all individual ORFs under the control of the strong constitutive *TEF2* promoter using synthetic genetic array technology. Double mutants were plated onto specific 96-well plates suited for fluorescence microscopy analysis, and individual wells were automatically imaged three times using high-throughput fluorescence microscopy. Automated thresholding identified mutants that showed significantly reduced relative GFP signal, indicative of enhanced reporter degradation (hits). (**E)** Summary of hits. 213 genes were identified in the first screen, 36 of which are annotated for encoding ER proteins, including members of the Hrd1 complex (left). 153 genes were identified in the second screen, 10 of which are annotated for encoding ER proteins. *PBN1* scored on both screens. See also suppl. Figure S1 and suppl. Table S1.

*Pbn1 shares structural homology with PIG-X and affects the turnover of misfolded Gas1 species with and without a GPI anchor* Pbn1 was initially reported to have an ER chaperone function, though its mechanisms of action remained unknown (Naik and Jones, 1998; Subramanian et al., 2006). Complementation cloning in mammalian cells identified Pbn1 as the functional homolog of mammalian PIG-X, involved in the transfer of mannose from dolichol phosphate-mannose to GlcN-(acyl)PI in conjunction with its catalytic subunit PIG-M (Gpi14 in yeast) during GPI anchor precursor synthesis, linking Pbn1 to ER-resident mannosyltransferases (Ashida et al., 2005). When we superimposed structural predictions of Pbn1 and its human homolog PIG-X, we found a common C-terminal TMD, an evolutionarily conserved domain proximal to the ER membrane, and an additional N-terminal domain present only in Pbn1, showing structural similarities between Pbn1 and PIG-X (Fig. 2A). These findings support that Pbn1 is part of the GPI-mannosyltransferase I (GPI-MT I) complex in yeast, likely explaining its essentiality. Using GFP-tagged Pbn1 for live cell microscopy, we found it co-localized with ER markers, whether expressed from its endogenous promoter (Fig. 2B) or upon overexpression from the *TEF2* promoter used in our screen (suppl. Fig. S2A).

**Figure 2.**
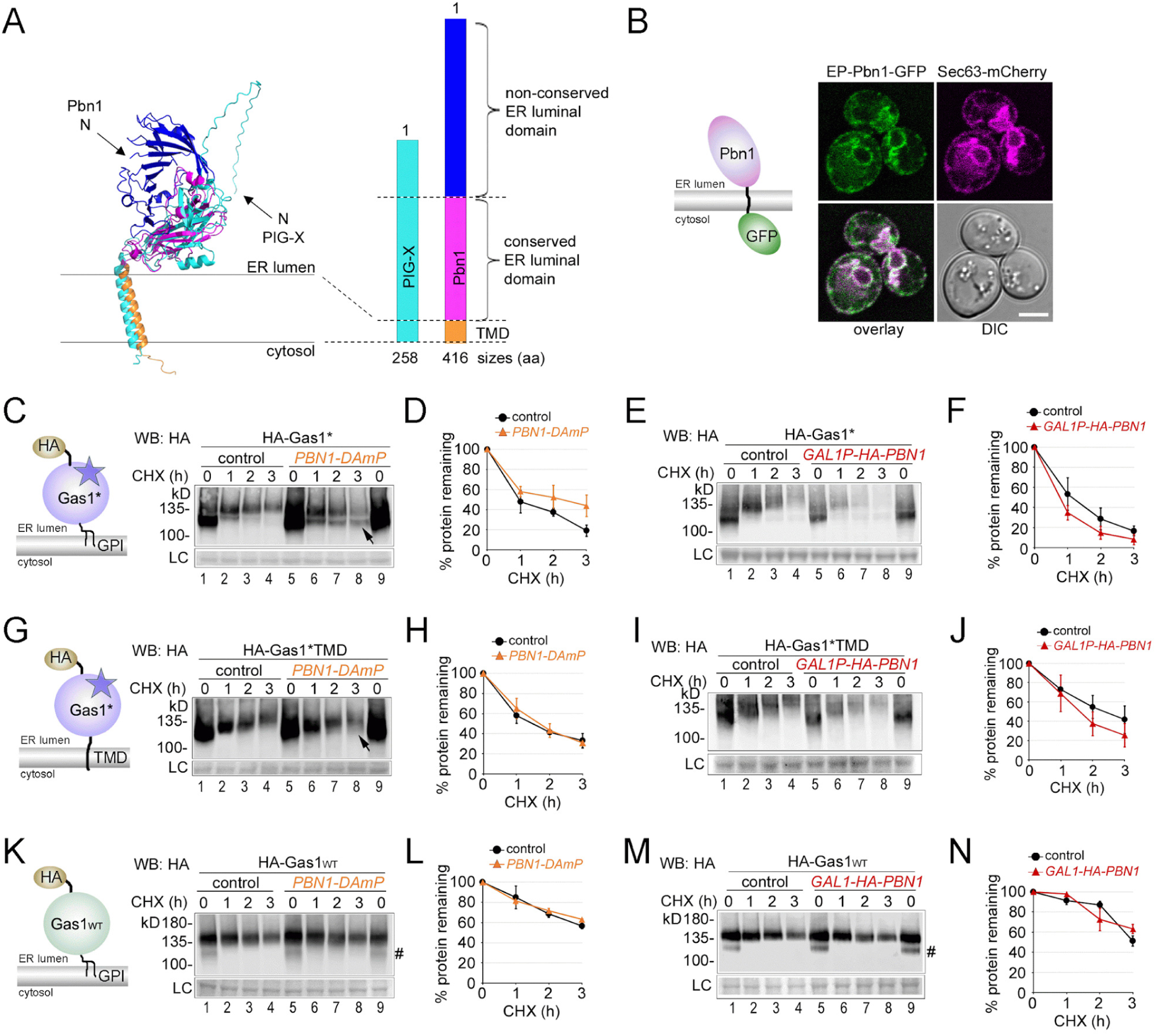
Pbn1 shares structural homology with PIG-X and affects the turnover of misfolded Gas1 species with and without a GPI anchor. **(A)** Alphafold structure predictions for yeast Pbn1 (blue/pink/orange) and human PIG-X (cyan) were superimposed using PyMOL (left panel). Protein lengths, their conserved and non-conserved domains, and their C-terminal TMDs are indicated with proportional bars (right panel). **(B)** Yeast cells co-expressing C-terminally GFP-tagged Pbn1 from its genomic locus and endogenous promoter (graphical depiction) and the ER marker Sec63-mCherry were analyzed by live cell fluorescence microscopy and differential interference contrast (DIC) microscopy. Scale bar: 3μm. **(C)** Control cells and cells containing the *PBN1-DAmP* allele were used to express plasmid-borne HA-tagged Gas1* (as depicted) for cycloheximide (CHX) shut-off experiments. Cells were lysed at indicated time points after application of CHX, and remaining HA-Gas1* was measured by SDS-PAGE and Western Blotting (WB) with antibodies against HA. Membrane staining with Ponceau served as loading control (LC). The arrow indicates a subpopulation of HA-Gas1* with lower MW in *PBN1-DAmP* cells compared to control cells. **(D)** Quantifications of results from experiments shown in (C). Total remaining protein (upper and lower bands combined) has been quantified. Mean values and standard deviations from at least three individual experiments are shown. **(E)-(F)** Control cells and cells that expressed genomically HA-tagged Pbn1 from the *GAL1* promoter (*GAL1P-HA-PBN1*) were used to express plasmid-borne HA-Gas1*. Experimental procedures and quantifications are described in (C) and (D) with the exception that cells were grown in a medium supplemented with galactose as carbon source for the expression of HA-Pbn1. **(G)-(J)**. As in (C)-(F), but expressing plasmid-borne HA- Gas1*TMD. HA-Gas1*TMD contains the N428Q mutation, which prevents TMD removal and GPI anchor attachment, thereby rendering HA-Gas1*TMD a TM protein. **(K)-(N)** As in (C)-(F), but expressing plasmid-borne HA-tagged wild-type Gas1 (HA-Gas1WT). Species with lower MW at times zero (#) represent the non-glycosylated ER form of HA-Gas1 (Sikorska et al., 2016). See also suppl. Figure S2.

To investigate the role of Pbn1 in the degradation of Gas1* *in vivo*, we measured protein degradation after blocking synthesis with cycloheximide (CHX) in both control cells and cells expressing the hypomorphic *PBN1-DAmP* allele, combined with Western Blot analysis. Degradation of Gas1* was preceded by an increase in its molecular weight (MW) (Fig. 2C, lanes 1-4), attributed to protein O-mannosylation by the Pmt1/2 complex, supporting previous data (suppl. Fig. S2B; and (Fujita et al., 2006; Goder and Melero, 2011)). We confirmed that deletion of the Pmt1/2 complex accelerated Gas1* degradation (suppl. Figs. S2C and D), due to a loss of ER retention and an increase in the fraction of the protein rapidly directed to the vacuole (Hirayama et al., 2008; Goder and Melero, 2011; Sikorska et al., 2016). Gas1* O- mannosylation occurs within the ER, rather than in the Golgi or other downstream organelles, as genetically blocking ER export did not reduce or eliminate the observed increase in MW of Gas1* (suppl. Figs. S2E and F).

In *PBN1-DAmP* cells, the degradation rate of Gas1* was reduced, and a subpopulation of the misfolded protein showed no increase in MW (Figs. 2C, lanes 5-8, arrow, and 2D). Conversely, overexpression of *PBN1* enhanced Gas1* degradation (Figs. 2E and F, and suppl. Fig. S2G). When expressing Gas1*TMD, a Gas1 mutant that fails to attach to a GPI anchor and retains its C-terminal TMD (Sikorska et al., 2016), we did not observe protein stabilization in PBN1-DAmP cells (Figs. 2G and H). However, we consistently observed a small subpopulation without an increase in MW, which was not detected in control cells (Fig. 2G, lane 8, arrow). Notably, similar to Gas1*, we observed faster degradation of Gas1*TMD in cells overexpressing *PBN1* (Figs. 2I and J). In contrast, the slower turnover rate and MW of HA-tagged wild-type Gas1 remained unchanged in both *PBN1-DAmP* cells and cells overexpressing *PBN1* (Figs. 2K-N). Collectively, these data demonstrate that Pbn1 influences the degradation of misfolded Gas1, regardless of the presence of a GPI anchor.

The changes in molecular weight (MW) of reporters in *PBN1-DAmP* cells suggested alterations in protein O-mannosylation. To investigate whether *PBN1-DAmP* cells indirectly affect protein O-mannosylation by altering PMT activity due to changes in GPI metabolism and ER homeostasis (Wang et al., 2020), we expressed Gas1* in the deletion mutants *Δted1* and *Δcwh43*. These mutants affect GPI metabolism by blocking GPI glycan and lipid remodeling, respectively, causing a strong unfolded protein response (UPR) (Jonikas et al., 2009). Protein O-mannosylation of Gas1* by the Pmt1/2 complex was unaffected in both mutants (suppl. Fig. S2H). Additionally, wild-type Gas1, which is O-mannosylated by Pmt4, displayed unaltered MW in *PBN1-DAmP* cells, indicating that Pmt4 activity was not affected (Fig. 2K). Together, these data suggest that Pbn1 may have a previously unknown role in the O-mannosylation of proteins, in addition to its function in the O-mannosylation of GPI anchor precursors.

### Pbn1 function is connected to protein O-mannosylation and ERAD

To investigate the potential role of Pbn1 in protein O-mannosylation, we used the well-studied model substrate CPY* along with a modified version. CPY* is typically degraded independently of O-mannosylation. Consistently, it displayed uniform MW during the CHX chase period, and its degradation rate was unaffected in both *PBN1-DAmP* cells and cells overexpressing *PBN1* (Figs. 3A-D). Next, we generated the modified version CPY*TMD by fusing the C-terminal domain of Gas1*TMD, which includes its TMD and part of its proximal SRR, to the C-terminus of CPY* (Fig. 3E, highlighted in yellow). The SRR of Gas1 is known to be the primary target for O-mannosylation (Nuoffer et al., 1993). In contrast to CPY*, CPY*TMD displayed an increase in MW prior to degradation (Fig. 3F, lanes 1-4). The increase in MW was due to O-mannosylation by the Pmt1/2 complex (suppl. Figs. S3A and B). Deletion of the Pmt1/2 complex not only caused a loss of O-mannosylation but also increased the vacuolar degradation of CPY*TMD, reflecting Gas1 phenotypes (suppl. Figs. S3A-D). In *PBN1-DAmP* cells, a fraction of CPY*TMD showed no increase in MW (Fig. 3F, lanes 5-9, arrow). Moreover, whereas the degradation rate of CPY*TMD was slightly reduced in *PBN1-DAmP* cells, it was accelerated upon *PBN1* overexpression, similar to results obtained with Gas1* and Gas1*TMD (Figs. 3F-I). Since the C-terminal domain from Gas1*TMD attached to CPY* is devoid of sites for N-glycosylation, the observed phenotypes with CPY*TMD ruled out alterations of mannoses on N-glycans and suggests that Pbn1, like Pmt1 and Pmt2, affects protein O-mannosylation.

**Figure 3.**
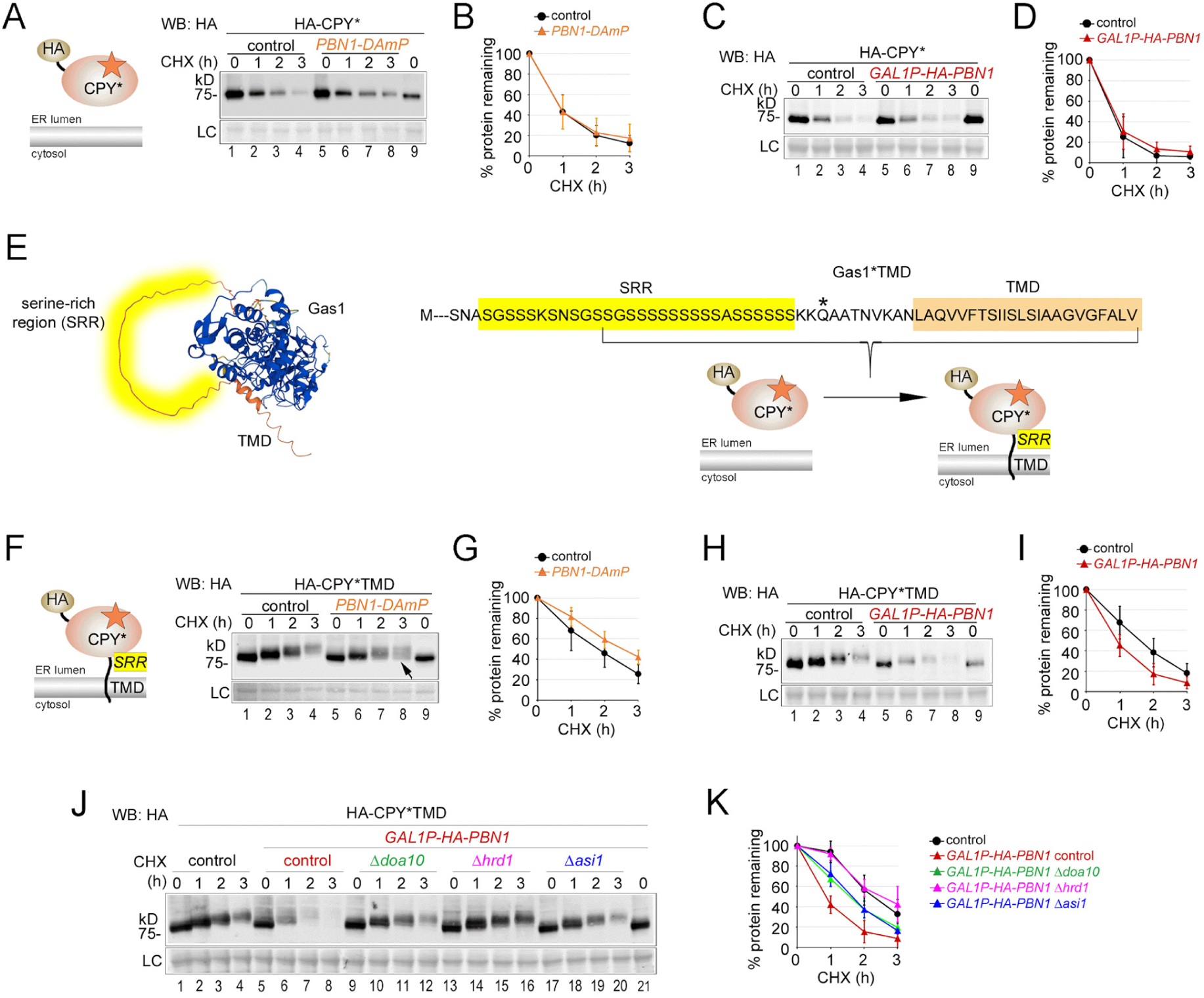
Pbn1 function is connected to protein O-mannosylation and ERAD. **(A)** Control cells and cells containing the *PBN1-DAmP* allele were used to express plasmid-borne HA-tagged CPY* (depiction) for CHX shut- off experiments. **(B)** Quantifications of results from experiments shown in (A). Mean values and standard deviations from at least three individual experiments are shown. **(C)-(D)** Control cells and cells containing *GAL1P-HA-PBN1* were used to express plasmid-borne HA-CPY* for CHX shut-off experiments in medium supplemented with galactose as carbon source for the expression of HA-Pbn1. **(E)** Left panel: Alphafold structure prediction of wild- type Gas1 with the C-terminal unstructured serine-rich region (SRR) (yellow) upstream of the TMD (orange). Right panel: The C-terminal region of Gas1*TMD is shown in single letter amino acid code with the SRR (Ser497-Ser525) and the TMD. The N428Q mutation that prevents GPI anchor attachment is marked with an asterisk (*). To generate HA-CPY*TMD, the indicated sequence of Gas1*TMD (in brackets) was fused with the C-terminus of HA-CPY*. **(F)-(I)** As in (A)-(D), but with plasmid-borne HA-CPY*TMD used for expression in the indicated control and mutant strains. **(J)-(K)** As in (C)-(D), but with plasmid-borne HA-CPY*TMD used for expression in control cells and in mutants lacking the indicated ERAD components. See also suppl. Figure S3.

To determine whether the increased degradation rate of CPY*TMD upon Pbn1 overexpression is linked to ERAD, we repeated the experiments in specific mutant backgrounds. Yeast contains three distinct ERAD complexes: the Hrd1-complex, the Doa10- complex, and the Asi-complex; for review, see (Krshnan et al., 2022). Deletion of any of the ERAD complexes reduced the degradation rate, with the strongest reduction observed in *Δhrd1* cells (Figs. 3J and K). In *Δemp24* cells, which are proficient in ERAD but deficient in vesicular ER export, CPY*TMD degradation was comparable to that in control cells (suppl. Figs. S3E and F).

Together, these data connect Pbn1 function to protein O-mannosylation and ERAD.

### Pbn1 is associated with Gpi14 in vivo

Complementation assays predicted Pbn1 to be functionally connected to the ER mannosyltransferase and GT-C superfamily member protein Gpi14 for GPI anchor biosynthesis (Ashida et al., 2005). To investigate whether Pbn1 is associated with Gpi14 *in vivo*, we utilized the observation that N-terminal tagging of Pbn1 with GFP caused its accumulation in ER puncta, independent of its expression levels (suppl. Figs. S4A and B). Whereas Gpi14-tdimer showed uniform expression across the perinuclear and peripheral ER, its co-expression with GFP-Pbn1 led to the nearly complete co-localization of the visible cellular pool of Gpi14 with Pbn1 in ER puncta (Figs. 4A and B). Co-expression of the utilized tagged versions of these essential proteins did not compromise their main functions, as no growth defects were observed under these conditions, even in the presence of the N-glycosylation inhibitor and ER stress inducer tunicamycin (suppl. Fig. S4C). These data demonstrate that Pbn1 is associated with Gpi14 *in vivo*, suggesting that Gpi14 may participate in protein O-mannosylation linked to ERAD.

**Figure 4.**
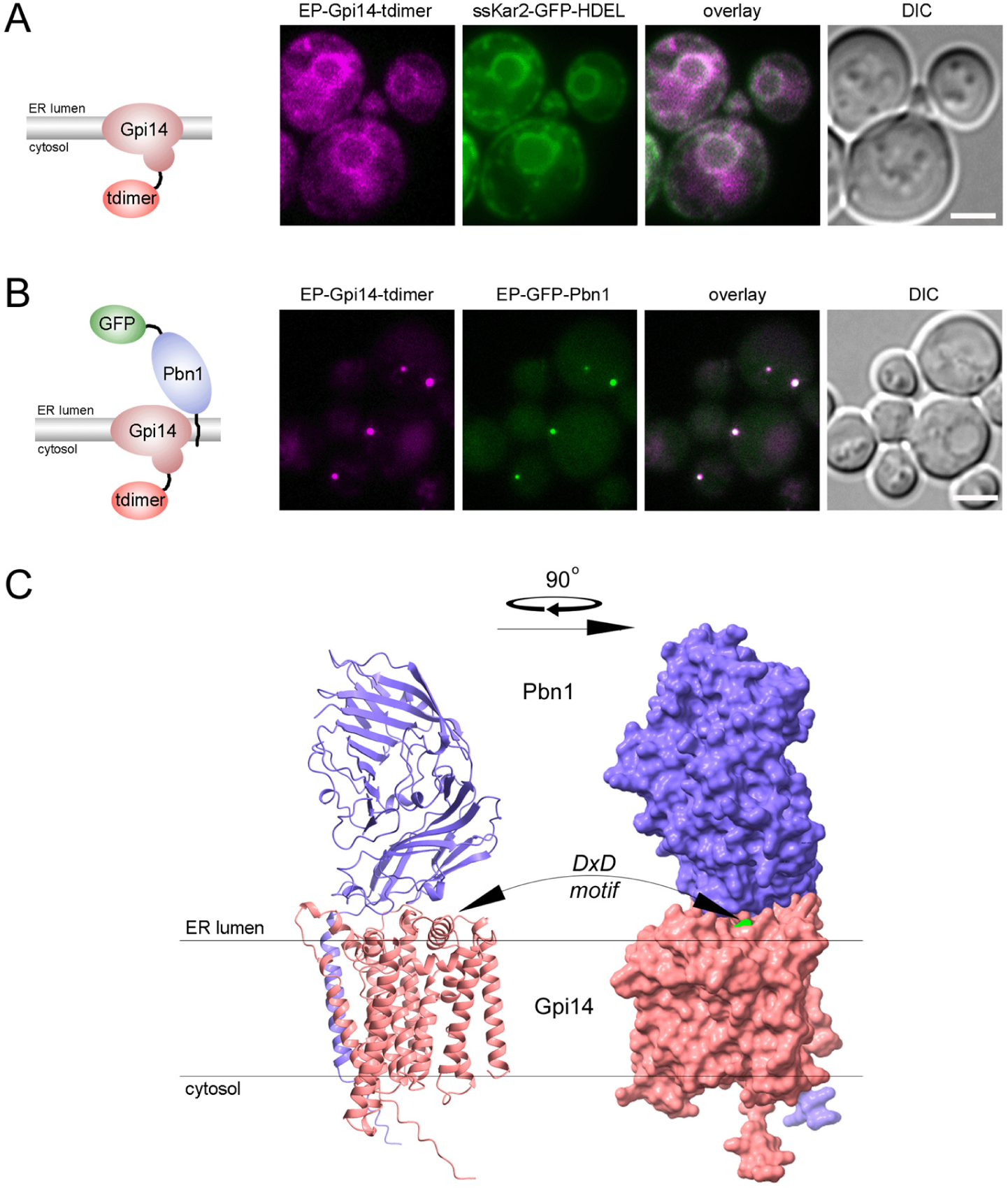
Pbn1 is associated with Gpi14 *in vivo*. **(A)** Yeast cells co-expressing C-terminally tdimer2-tagged genomic Gpi14 (as depicted ) and the ER marker ssKar2-GFP-HDEL were analyzed by live cell fluorescence microscopy and DIC microscopy. Scale bar: 2μm **(B)** Yeast cells co-expressing C-terminally tdimer2-tagged genomic Gpi14 and N-terminally GFP-tagged genomic Pbn1 (depiction) were analyzed by live cell confocal fluorescence microscopy and DIC microscopy. Scale bar: 3μm. **(C)** Alphafold multimer prediction for a heterodimer containing Pbn1 (purple) and Gpi14 (pink). The catalytic DxD motif (green) in Gpi14 is localized within a cleft near the luminal side of the ER membrane oriented toward the conserved luminal domain of Pbn1. See also suppl. Figure S4.

The AlphaFold multimer structure of the simplest stoichiometric complex between Pbn1 and Gpi14, a heterodimer, provides a potential explanation for how the non-catalytic Pbn1 might regulate O-mannosylation across a range of substrates (Fig. 4C). Pbn1 forms part of an interface together with Gpi14 that is primed to regulate substrate access to the conserved catalytic DxD motif in Gpi14 (Fig. 4C, motif depicted in green, and (Wiggins and Munro, 1998)).

### Pbn1-Gpi14 protects cells from ER stress caused by inhibition of N-glycosylation and promotes O-mannosylation and ERAD of a misfolded non-N-glycosylated model protein

Our findings suggest that Pbn1 participates in generating ERAD cues through protein O-mannosylation on a set of misfolded proteins. These misfolded proteins contain an SRR, which is a preferred target for protein O-mannosylation, and N-glycans, which can generate distinct and potent ERAD cues. Next, we aimed to determine whether Gpi14 participates in protein O-mannosylation and whether the Pbn1-Gpi14 complex has physiological relevance in the context of ERAD, including for proteins that lack N-glycans and SRRs. To this end, we individually tagged Pbn1 and Gpi14 at their genomic loci with the auxin-inducible degron AID* in combination with GFP. In a functionality test, we confirmed that Pbn1-AID*GFP was degraded and no longer detectable by Western blot analysis after the application of auxin, providing us with a tool to rapidly deplete Pbn1 or Gpi14 for phenotypic analyses (suppl. Fig. S5A).

First, we compared the growth rates of control cells with those of cells expressing Pbn1 or Gpi14 tagged with the degron in the absence or presence of auxin and tunicamycin. Tunicamycin generates a large pool of newly synthesized non-N-glycosylated ER proteins with a tendency to misfold (Fig. 5A). Cellular growth in rich medium was unaffected in cells expressing degron-tagged Pbn1 or Gpi14, indicating that the tagged proteins were functional (Fig. 5A, YPD). As expected for the depletion of essential genes, cells expressing degron-tagged Pbn1 or Gpi14 showed reduced growth after the application of auxin (Fig. 5A, YPD+IAA). In the presence of tunicamycin, all cells displayed growth defects compared to control medium, with the strongest effect observed in the control strain *Δire1*, which was unable to mount the UPR (Fig. 5A, YPD+Tm). However, control cells and cells expressing degron-tagged Pbn1 or Gpi14 showed similar growth rates. In the presence of tunicamycin combined with auxin, cells expressing Pbn1-AID*GFP and Gpi14-AID*GFP displayed a stronger growth defect than control cells (Fig. 5A, YPD+IAA+Tm). These results indicate that Pbn1-Gpi14 can alleviate ER stress caused by tunicamycin. Together with our previous data, this suggests that Pbn1-Gpi14 has physiological relevance in the context of ERAD for insufficiently or non-N-glycosylated misfolded proteins, including those that lack SRRs.

**Figure 5.**
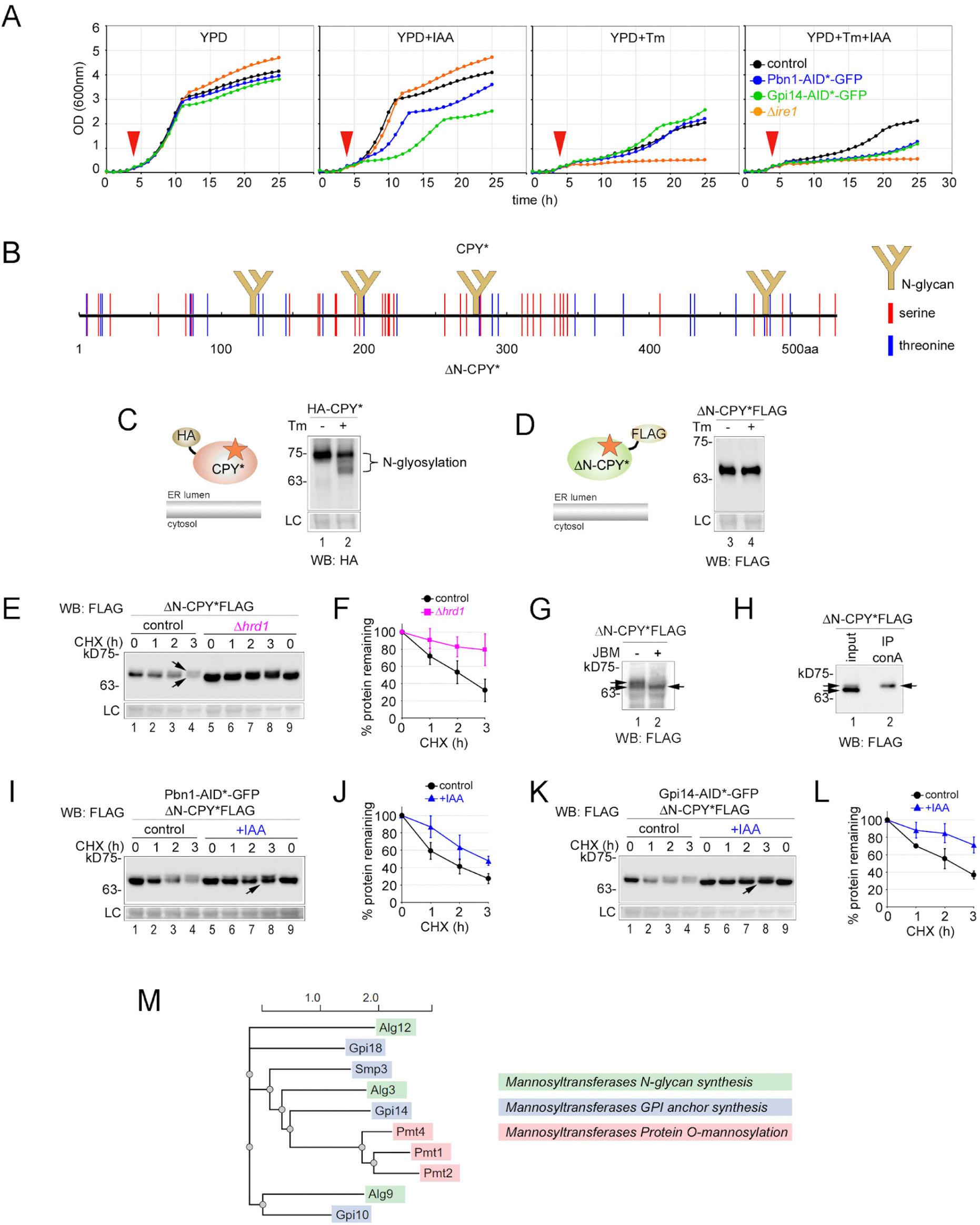
Pbn1-Gpi14 protects cells from ER stress caused by inhibition of N-glycosylation and promotes O-mannosylation and ERAD of a misfolded non-N-glycosylated model protein. **(A)** The indicated strains were grown in rich medium (YPD) in absence or presence of 1μg/ml tunicamycin (Tm) and 500μM auxin (indole-3-acetic acid (IAA)) at 30°C. Drugs together with solvents or solvents alone were applied four hours after starting the experiment by resetting cellular density (indicated by red arrowheads). Growth was determined by automated measurement of absorbance at 600 nm for individual cell cultures over a period of 25 hours. The values shown are the mean results from two independent measurements. **(B)** Schematic illustrating the properties of CPY* and ΔN- CPY* protein sequences. CPY* is modified with four N-linked glycans at the indicated positions, whereas ΔN-CPY* contains mutated N-glycosylation sites and lacks N-glycans. Both proteins contain identical serines and threonines distributed throughout their sequences, as indicated. However, both lack the serine-rich region present in CPY*TMD (see suppl. Fig. 5B). **(C)** Lysates from cells expressing HA-tagged CPY* were analyzed by Western Blot (WB) analysis before and after cells were grown in the presence of 1 μg/ml tunicamycin (Tm) for one hour. **(D)** Like in (C) using cells expressing FLAG-tagged ΔN-CPY*. **(E)** Control and Δ*hrd1* cells were used to express plasmid-borne ΔN-CPY*FLAG (graphical depiction) for CHX shut-off experiments. The arrows indicate the presence of at least two protein populations with different MWs. **(F)** Quantifications of results from experiments shown in (E). Mean values and standard deviations from at least three individual experiments are shown. **(G)** Cells expressing ΔN- CPY*FLAG were lysed, and the lysate was incubated either with jack-bean-mannosidase (+) or with solvent (-) for four hours, followed by WB analysis. The arrow indicates the visible band with higher MW that was sensitive to mannosidase treatment. **(H)** Cells expressing ΔN-CPY*FLAG were lysed, and the lysate was incubated with concanavalin A(conA)-sepharose for three hours, followed by precipitation and WB analysis. The arrows indicate that the visible band with higher MW was precipitated with conA beads. **(I)** Cells expressing genomic Pbn1 C- terminally tagged with the degron AID* in combination with GFP were used to express plasmid-borne ΔN- CPY*FLAG for CHX shut-off experiments. Three hours before applying CHX, cells were supplemented with auxin (+IAA) or only with solvent (control). The arrow indicates the accumulation of the population of ΔN-CPY*FLAG with lower MW. **(J)** Quantifications of results from experiments shown in (I). Mean values and standard deviations from at least three individual experiments are shown. **(K)-(L)** As in (I)-(J), but using cells that expressed genomic Gpi14 tagged with the degron. **(M)** Dendrogram from a phylogenetic analysis of the amino acid sequences of the indicated members of the ER-resident GT-C superfamily of O-mannosyltransferases, using NGPhylogeny.fr (Lemoine et al., 2019). The color bars refer to annotated enzyme functions. See also suppl. Figure S5.

To directly address this, we analyzed the role of Pbn1-Gpi14 in the degradation of a misfolded, non-N-glycosylated ER protein under non-stressed conditions. We used FLAG-tagged ΔN-CPY*, which, unlike CPY*, has all its four N-glycosylation sites removed (Fig. 5B; Kostova and Wolf, 2005). Additionally, unlike CPY*TMD, ΔN-CPY* does not contain an SRR (Fig. 5B; suppl. Fig. S5B). However, all CPY* versions contain a multitude of serines and threonines scattered throughout their sequences (Fig. 5B; suppl. Fig. S5B). Whereas N- glycosylated CPY* was sensitive to tunicamycin, ΔN-CPY* was insensitive to the drug, confirming the lack of N-glycosylation sites (Figs. 5C and D). ΔN-CPY* was degraded via Hrd1- dependent ERAD, albeit less efficiently than CPY*, demonstrating that it was not aggregated and remained retrotranslocation-competent (Figs. 5E and F). Unlike CPY*, ΔN-CPY* displayed a subpopulation with higher MW (Fig. 5E, lane 4, arrows; compare with Fig. 3A, lane 4). After treatment of cell lysate with jack-bean mannosidase, the subpopulation with higher MW was no longer detectable (Fig. 5G, lane 2), showing that ΔN-CPY* was O-mannosylated. Supporting this, we could specifically immunoprecipitate the ΔN-CPY* subpopulation with higher MW using the lectin concanavalin A, which only binds glycosylated proteins. ΔN-CPY* with lower MW and the non-glycosylated control protein CFTR failed to be precipitated (Fig. 5H; suppl. Fig. S5D).

Next, we measured the degradation rates of ΔN-CPY* in cells expressing Pbn1-AID*GFP with and without auxin-induced degradation. Depletion of Pbn1 resulted in a reduced degradation rate of ΔN-CPY* (Figs. 5I and J). At the same time, the subpopulation of ΔN-CPY* with lower MW accumulated, indicating deficient O-mannosylation (Fig. 5I, compare lanes 4 and 8, arrow). In contrast to ΔN-CPY*, the degradation of N-glycosylated CPY* was unaffected by Pbn1 depletion, consistent with data obtained from *PBN1-DAmP* cells (suppl. Figs. S5E and F, and Fig. 3A). In addition to the unchanged MW through the chase period, this further supports that O-mannosylation of misfolded CPY* occurs only in the absence of N-glycans. Similar to the effects observed with Pbn1, the depletion of Gpi14 resulted in the stabilization of ΔN-CPY* and the accumulation of its non-O-mannosylated form (Figs. 5K, compare lanes 4 and 8, arrow, and L). Additionally, ΔN-CPY* degradation was stabilized and the non-O-mannosylated form accumulated in cells containing either the *PBN1-DAmP* or *GPI14-DAmP* allele (suppl. Figs. S5G and H). Interestingly, we did not detect a contribution from canonical PMTs to the O-mannosylation and ERAD of ΔN-CPY* (suppl. Figs. S5I and J). This specificity highlights the observations made with Pbn1 and Gpi14-depleted cells and suggests that Pbn1- Gpi14 functions as a major protein O-mannosyltransferase for ΔN-CPY*, a misfolded protein that lacks N-glycans and SRRs, thereby promoting its degradation.

Given our functional data linking Pbn1-Gpi14 to protein O-mannosylation, we conducted a phylogenetic analysis to assess the evolutionary distance of Gpi14 to canonical PMTs. Based on the amino acid sequences of all major ER-resident O-mannosyltransferases of the GT-C superfamily, Gpi14 is positioned closest to the PMTs (Fig. 5M). These findings suggest that protein O-mannosylation in yeast predates the emergence of canonical PMTs in evolution.

## Discussion

Our results revealed that protein O-mannosylation can generate ERAD cues. Protein O-mannosylation has been previously shown to terminate futile folding cycles, maintain protein solubility, and promote ER retention, all of which are prerequisites for protein retrotranslocation (Harty et al., 2001; Nakatsukasa et al., 2004; Hirayama et al., 2008; Goder and Melero, 2011; Xu et al., 2013). Collectively, these data show that protein O-mannosylation is a key modification in various steps of ERQC and ERAD.

Although both O-mannosylation- and N-glycosylation-dependent ERAD cues are glycan-based, they fundamentally differ in their production. N-linked ERAD cues are generated by trimming pre-attached N-glycans, and their number and location on a given protein are fixed. In contrast, O-linked ERAD cues are generated *de novo*, and on demand, with much greater flexibility regarding their location on the protein, as they can be constructed on virtually any accessible hydroxyl-amino acid. Our data suggest that O-linked ERAD cues add a layer of control for the degradation of N-glycosylated proteins and initiate a fail-safe mechanism for degrading misfolded proteins when other ERAD cues are insufficient or absent.

Our findings related to SRRs support the first scenario. SRRs are hotspots for protein O-mannosylation by PMTs and are found in proteins that are usually N-glycosylated as well, such as many GPI-APs in fungi (de Groot et al., 2003; Lommel and Strahl, 2009). SRRs may be beneficial for specific substrates, and our data suggest that they provide autonomous platforms that can connect ERQC with ERAD, even in the presence of separately processed N-glycans. For instance, in the case of Gas1, an endogenous GPI-AP in yeast, only the misfolded version Gas1*, but not the wild-type protein, becomes O-mannosylated on its SRR primarily by the Pmt1/2 complex and to a minor degree by Pbn1-Gpi14. This modification leads to ER retention and thus prolonged time for ER-assisted folding (Hirayama et al., 2008; Goder and Melero, 2011; Sikorska et al., 2016). Our data obtained with the SRR-containing fusion protein CPY*TMD show that protein O-mannosylation of an SSR can be extensive, and increasing O-mannosylation may be a mechanism to shift a protein from ER retention and folding to ERAD. With a single mannose having a molecular weight (MW) of 180 Da, the observed increase in MW of CPY*TMD by more than 10 kDa suggests that protein O-mannosylation can include the formation of structures with multiple glycans. Whether these structures are linear or branched mannose chains or mixed glycans built on top of single mannoses remains to be investigated. A similar scheme of signal build-up related to protein degradation is found in the ubiquitin space, where heterotypic and branched ubiquitin chains regulate degradation efficiency (Meyer and Rape, 2014; Leto et al., 2019). The receptor for O-linked ERAD cues is still unknown. It is reasonable to assume that it is a lectin, with the primary candidate being Yos9 (OS9 in mammals), which recognizes α1,6-linked terminal mannoses on trimmed N-glycans (Quan et al., 2008).

The second scenario is supported by our findings related to CPY* and the non-N-glycosylated version ΔN-CPY*. Protein O-mannosylation of ΔN-CPY* by Pbn1-Gpi14, but not of CPY*, demonstrates that *de novo* O-mannosylation can function as a fail-safe mechanism for promoting degradation in the absence of other ERAD cues. The slower degradation kinetics of ΔN-CPY* compared to CPY* suggests that O-glycan-dependent ERAD operates with slower kinetics than N-glycan-dependent ERAD. Interestingly, PMTs do not appear to participate in the degradation of ΔN-CPY*. While we cannot formally exclude redundancies of the remaining PMTs in the utilized deletion mutants, scattered hydroxyl-amino acids may not be primary targets for O-mannosylation by PMTs, at least under unstressed conditions. Indeed, besides serine-rich domains, the motifs for PMT-dependent O-mannosylation appear complex and might be structure- rather than sequence-based (Lommel and Strahl, 2009). Since the UPR includes the upregulation of PMTs and an ensuing increase in global protein O-mannosylation, a possible scenario predicts changes in substrate accessibility or PMT processivity under these conditions.

Unlike canonical PMTs, Pbn1-Gpi14 was not upregulated as part of the UPR (Travers et al., 2000). This might not be surprising given that bifunctional Pbn1-Gpi14 has roles in GPI anchor biosynthesis and protein O-mannosylation. GPI metabolism and ER membrane homeostasis are tightly connected, and upregulation of GPI biosynthesis exacerbates rather than alleviates ER stress (Wang et al., 2020). Accordingly, GPI-APs are rapidly escorted out of the ER under stress conditions, probably to maintain ER membrane homeostasis (Satpute-Krishnan et al., 2014). This, in combination with our data from growth assays with degron-tagged Pbn1 and Gpi14, suggests that Pbn1-Gpi14 maintains ER homeostasis through the regulation and promotion of fail-safe ERAD of a range of cellular substrates, rather than reestablishing ER homeostasis after ER stress.

Pbn1-Gpi14 was first linked to the transfer of mannose to GlcN-(acyl)PI for GPI anchor precursor biosynthesis (Ashida et al., 2005). Although the transfer of mannoses to either GlcN- (acyl)PI or hydroxyl-amino acids involves comparable chemistry, the binding of different substrates predict different Pbn1-Gpi14 conformations or associations with various cellular factors, including the formation of different stoichiometric complexes. Interestingly, Gpi14 overexpression alone affects conjugation of mannose to the GPI precursor in the parasite *Leishmania*; thus, catalytic Gpi14 can act without Pbn1 for GPI precursor synthesis (Ribeiro et al., 2019). The structural model of Pbn1-Gpi14 supports flexibility and shows similarities to protein O-mannosyltranferases. Pbn1 provides a large luminal domain to Gpi14, similar to the MIR domain that grants large luminal domains to PMTs. The distantly related transmembrane and tetratricopeptide repeat-containing (TMTC) protein O-mannosyltransferase family also possesses a large luminal domain made up of tetratricopeptide repeats (TPRs) (Larsen et al., 2017). MIR domains and TPRs have been implicated in protein and peptide binding (Larsen et al., 2017; Chiapparino et al., 2020). In contrast, luminal domains are smaller or absent in O-mannosyltransferases exclusively involved in the biosynthesis of N-glycans and GPI anchors, which typically possess fewer or exclusive substrates (Albuquerque-Wendt et al., 2019; Alexander and Locher, 2023).

## Supporting information

suppl figures Lemus et al

Table S1 Lemus et al

Table S2 Lemus et al

Table S3 Lemus et al

## Acknowledgements

We thank Enrique Alanis for help with figures, Mar Bustamante and Carlos Ruiz for help with experiments, Piet de Groot for advice, Yoshifumi Jigami, Tom Rapoport, Pedro Carvalho, and Jonathan Weissman for reagents, and Pedro Carvalho and Sabine Strahl for critical reading of the manuscript and for many helpful comments. M.S. was supported by grants of the EU-H2020-ERC-CoG (OnTarget 864068). The robotic system of the Schuldiner lab was purchased through the kind support of the Blythe Brenden-Mann Foundation. M.S. is an Incumbent of the Dr. Gilbert Omenn and Martha Darling Professorial Chair in Molecular Genetics. L.L. and V.G. were supported by a grant of the Agencia Estatal de Investigación (AEI/10.13039/501100011033/PID2022-136665NB-I00). L.L. was supported by the European Regional Development Fund from the Junta de Andalucía.

## Author contributions

L.L., A.R., M.S., and V.G. designed experiments, L.L., A.R., H.M., and V.G. conducted experiments, L.L., A.R., H.M., M.S. and V.G. evaluated data, L.L. and V.G. wrote the manuscript, L.L., M.S. and V.G. edited the manuscript.

## Material and Methods

### Yeast strains used in this study

All experiments were performed using common laboratory yeast strains constructed in a BY (MATa/α *his3Δ1/his3Δ1 leu2Δ0/leu2Δ0 LYS2/lys2Δ0 met15Δ0/MET15 ura3Δ0/ura3Δ0*) or W303 (*leu2-3,112 trp1-1 can1-100 ura3-1 ade2-1 his3-11,15*) background. Genomic tagging of yeast strains was performed using standard PCR-based amplifications from suitable plasmids followed by transformation with PCR products and PCR-based verifications and/or Western Blot analysis. A list of yeast strains used in this study is found in supplementary Table S2.

### Plasmids used in this study

A list of plasmids used in this study is found in supplementary Table S3. All plasmid-borne constructs were generated by standard PCR-based amplifications and/or restriction enzyme-based subcloning. All relevant constructs have been verified by sequencing.

### Genome-wide yeast screens

Screen using Leu2-TMD-Gas1* as a reporter. The Yeast Knock-out (YKO) deletion collection and the *DAmP* collection were grown to logarithmic growth phase in 96-well plates containing rich medium and transformed with plasmid VGp161, containing Leu2-TMD-Gas1*. Transformants were selected by growth in 96-well plates containing synthetic media lacking uracil. Transformants were transferred to 96-well plates containing synthetic media lacking uracil and in parallel to 96-well plates containing synthetic media lacking both uracil and leucine. Plates were incubated at 30°C and evaluated for growth after five days. Transformants with growth in both media were designated as hits and are listed in supplementary Table S1. Screen using GFP-Gas1* as a reporter: GFP-tagged. BY4741 containing a *Δted1* deletion was used for genomic integration of GFP-Gas1* (VGc226) for expression from its endogenous promoter. The strain was crossed with the collection of yeast strains expressing N-terminal mCherry fusions to all individual ORFs under the control of the strong constitutive *TEF2* promoter (Weill et al., 2018). For automated microscopy, cells were transferred from agar plates into 384-well polystyrene plates (Greiner) for growth in liquid media using the RoToR arrayer robot (Singer Instruments). Liquid cultures were grown in a shaking incubator (Liconic), overnight at 30°C. A Freedom EVO 200 liquid handler (Tecan) connected to the incubator was used to prepare the cells for microscopy by transferring them into glass-bottom 384-well microscope plates (Matrical Bioscience) coated with Concanavalin A (Sigma-Aldrich). After 20 min, cells were washed three times in complete medium to remove non-adherent cells and to obtain a cell monolayer. The plates were then transferred to an automated inverted fluorescent microscope system (Olympus) harboring a spinning disk module (Yokogawa CSU-W1 spinning disk confocal scanner with 50 μm pinhole) using a robotic swap arm. Images of cells in the 384-well plates were recorded in the same liquid as the washing step at 24°C using a 60× air lens (NA 0.9) and with a Hamamatsu ORCA-Flash 4.0 camera. Images of GFP (wavelength of 488 nm) were acquired following excitation by a laser and using an emission filter of 513/30 nm). Automated thresholding was performed to identify hits that showed >20% signal reduction compared to the median GFP signal. Hits are listed in supplementary Table S1.

### Plating assays

Cells were grown in the indicated liquid medium overnight at 30°C, adjusted to the same optical density and spotted with identical volumes in 1:10 dilution series onto plates containing synthetic medium lacking the indicated components. Plates were incubated for the indicated times and temperatures prior to imaging.

### Cycloheximide (CHX) shut-off experiments

The experiments were started with exponentially growing cells at 30°C in synthetic medium with an optical density (OD600) between 0.5 and 0.8. Translation was stopped by addition of CHX to a final concentration of 200μg/ml. Equal volume aliquots of cell culture were removed at indicated time points and moved to ice. Cells were pelleted, resuspended in 500μl cold 150mM NaOH and left on ice for five minutes, centrifuged and lysed by adding sample buffer containing 2% SDS and heating to 65°C for 10 min.

### Treatment of cells with tunicamycin

Tunicamycin (SIGMA) dissolved in DMSO was applied to exponentially growing cells to a final concentration of 1μg/ml, or to the concentration indicated. Control cells were treated with the same volume of DMSO (solvent). Cells were lysed after the indicated times and processed like non-treated cells for subsequent analyses.

### SDS-PAGE and Western Blotting

Samples were analyzed by standard SDS-PAGE followed by Western blotting using the indicated primary antibodies, peroxidase-coupled secondary antibodies (Roche) and ECL (PIERCE) as substrate. Images were taken with a LICOR imaging system and bands were quantified using Image Studio software (LICOR).

### Treatment with PNGaseF

Exponentially growing cells were lysed in cold phosphate buffered saline pH7.4 containing protease inhibitor cocktail (Roche) using a FastPrep homogenizer (MP Biomedicals) in combination with glass beads and bead beating. The lysate was transferred and SDS was added to a final concentration of 0.5%. Lysate was heated to 95°C for 10min, vortexed briefly and transferred to a new tube. Prior to the addition of PNGase F (New England Biolabs), G7 buffer and NP40 (both New England Biolabs) were added according to manufacturer’s instructions. Lysate supplemented with PNGase F and control lysate were incubated for 2 hours at 37°C. The reaction was terminated by the addition of sample buffer containing 2% SDS and heating to 65°C for 20 minutes.

### Treatment with α-Mannosidase from Canavalia ensiformis (Jack bean)

The reaction was performed according to the manufacturer’s instructions. In brief, cells were pelleted, washed and resuspended in sodium acetate buffer pH 4.5, together with 2mM ZnCl_2_ and protease inhibitor cocktail (Roche). Cells were lysed using glass beads and bead beating with a FastPrep homogenizer (MP Biomedicals). The lysate was transferred and SDS was added to a final concentration of 0.5%. Lysate was heated to 95°C for 10min, vortexed briefly and transferred to a new tube. NP-40 was added to a final concentration of 1%. Sample was split into two equal parts. 5μl of a 3M ammonium sulfate solution containing jack bean mannosidase (SIGMA) was added to one sample and the same amount of 3M ammonium sulfate to the control sample. Samples were incubated at 37°C for 4 hours, with stirring at 500 rpm. The reaction was terminated with the addition of sample buffer containing 2% SDS and heating to 65°C for 20 minutes.

### Precipitation of glycosylated protein with conA-Sepharose

Exponentially growing cells were pelleted, moved to ice and resuspended in 1 ml 150mM NaOH, left for 5 minutes on ice and pelleted. For lysis, cells were resuspended in 150 μl phosphate buffered saline pH 7.4, containing protease inhibitor cocktail (Roche), 2% SDS and 10mM dithiothreitol (DTT) and incubated at 95°C for 10min. Cells were briefly vortexed, pelleted and the supernatant was transferred to a new tube. An aliquot was removed for “input” and to the rest 1250 μl phosphate buffered saline pH 7.4, containing protease inhibitor cocktail (Roche), 100 μl washed concanavaline-Sepharaose beads (Amersham) and 1% bovine serum albumin (BSA) was added. The lysate was incubated rotating at 4°C for 3 hours. Beads were pelleted and washed four times with phosphate buffered saline pH 7.4, containing protease inhibitor cocktail (Roche) and 0.1% SDS. Bound material was eluted by incubation with sample buffer containing 2% SDS and heating to 65°C for 20 minutes.

### Growth assays

Cells were grown in the indicated liquid medium overnight at 30°C, reset to an optical density (OD600nm) of 0.05 in a volume of 500μl in a 24-well plate in duplicates and incubated for 24 hours in a BMG FLUOstar Omega multi-mode microplate reader at 700 rpm and a temperature of 30°C. Indicated drugs or only solvents were applied at the indicated times. ODs were recorded automatically every hour.

### GFP-processing assays

Cells were grown overnight, diluted to OD 0.2 and regrown for 5 hours. Removed aliquots were lysed by alkaline treatment (Kushnirov, 2000), resuspended in a cell-density normalized volume of loading buffer, followed by SDS-PAGE and Western Blotting using anti-GFP antibody (Roche), HRP-conjugated anti-mouse secondary antibody (Roche) and ECL (PIERCE) as substrate. Images were taken with a LAS-3000 mini imaging system (Fujifilm) and bands were quantified using Multi-Gauge software (Fujifilm).

### Fluorescence microscopy

Exponentially growing cells were washed with PBS and immediately analyzed by fluorescence microcopy. Cells were observed with a LEICA DMi8 microscope equipped with a 100x/1.4 oil Plan-Apo immersion lens and a DIC prism and polarizer for Normarksi imaging. Images were acquired using a Hamamatsu C1 3440-20CV camera and the LASX controller software (Leica).

### Confocal live cell microscopy

Exponentially growing cells were washed with PBS and scanned with a Laser Scanning Confocal Microscope from Zeiss (LSM 7 Duo) equipped with a BiG (binary GaAsP) module using a Plan-Apochromat objective 63x/1,40 Oil DIC. A 488nm Argon Laser (GFP) and a 561nm Helium-Neon Laser were used (tdimer, mCherry).

### Structure predictions

Superpositioning of Alphafold structure predictions was done with PyMOL using the align function. The superimposed structures were visualized using PyMOL’s graphical user interface. Protein complex prediction was done with AlphaFold Multimer in combination with ChimeraX (version 1.7). The protein complex was visualized in ChimeraX using a combination of ribbon and surface representations. The individual proteins were colored distinctly for clarity. Specific residues involved in catalytic activity were highlighted and labeled.

### Phylogenetic analysis

The analysis was performed according to instructions (https://ngphylogeny.fr/). All amino acid sequences were provided in FASTA format and the fully automated workflow with PhylML as tree inference was selected. This included the following steps and default parameters: Multiple Alignment (MAFFT: flavour: auto; Gap extension penalty: 0.123; Gap opening penalty: 1.53), Alignment Curation (BMGE: Sliding window size: 3; Maximum entropy threshold: 0.5; Gap rate cutoff: 0.5; Minimum block size: 5; Matrix: BLOSUM62), Tree inference (PhyML: Statistical criterion to select the model : AIC; Tree topology search : SPR; Branch support: No branch support; Proportion of invariant sites : Estimated; Number of categories for the discrete gamma model : 4; Parameters of the gamma model: estimated; Tree topology search: SPR; Optimise parameters: tlr; Model: LG; Equilibrium frequencies : ML model) and Tree Rendering (Newick Display: Model: LG; Equilibrium frequencies: estimated; Gamma distributed rates across sites: Yes; Gamma distribution parameter: 1.0; Remove gap strategy: Pairwise deletion of gaps; Starting tree: BioNJ; Tree Refinement: BalME SPR; Bootstrap: No; Decimal precision for branch lengths: 6).

## Abbreviations

ERQC: endoplasmic reticulum protein quality control
ERAD: endoplasmic reticulum- associated protein degradation
PMT: protein mannosyltransferase
TMTC: proteins: tetratricopeptide repeat-containing proteins
GPI: anchor: glycosylphosphatidylinositol anchor
TMD: transmembrane domain
GT-C: superfamily: glycosyltransferase C superfamily
GlcN-: (acyl)PI: glucosamine- (acyl)phosphatidylinositol
SRR: serine-rich region
UPR: unfolded protein response

## References

Albuquerque-Wendt, A., H.J. Hütte, F.F.R. Buettner, F.H. Routier, and H. Bakker. 2019. Membrane Topological Model of Glycosyltransferases of the GT-C Superfamily. Int. J. Mol. Sci. Artic. doi:10.3390/ijms20194842.

Alexander, J.A.N., and K.P. Locher. 2023. Emerging structural insights into C-type glycosyltransferases. Curr. Opin. Struct. Biol. 79:102547. doi:10.1016/J.SBI.2023.102547.

Ashida, H., Y. Hong, Y. Murakami, N. Shishioh, N. Sugimoto, Y.U. Kim, Y. Maeda, and T. Kinoshita. 2005. Mammalian PIG-X and yeast Pbn1p are the essential components of glycosylphosphatidylinositol-mannosyltransferase I. Mol. Biol. Cell. 16:1439–1448. doi:10.1091/MBC.E04-09-0802/ASSET/IMAGES/LARGE/ZMK0030530530007.JPEG.

Bai, L., A. Kovach, Q. You, A. Kenny, and H. Li. 2019. Structure of the eukaryotic protein O-mannosyltransferase Pmt1-Pmt2 complex. Nat. Struct. Mol. Biol. 26:704–711. doi:10.1038/S41594-019-0262-6.

Biederer, T., C. Volkwein, and T. Sommer. 1997. Role of Cue1p in ubiquitination and degradation at the ER surface. Science (80*-.).* 278:1806–1809.

Bloch, J.S., G. Pesciullesi, J. Boilevin, K. Nosol, R.N. Irobalieva, T. Darbre, M. Aebi, A.A. Kossiakoff, J.-L. Reymond, and K.P. Locher. 2020. Structure and mechanism of the ER-based glucosyltransferase ALG6. Nat. |. 579:443. doi:10.1038/s41586-020-2044-z.

Bodnar, N.O., and T.A. Rapoport. 2017. Molecular Mechanism of Substrate Processing by the Cdc48 ATPase Complex. Cell. 169. doi:10.1016/j.cell.2017.04.020.

Caramelo, J.J., O.A. Castro, G. De Prat-Gay, and A.J. Parodi. 2004. The Endoplasmic Reticulum Glucosyltransferase Recognizes Nearly Native Glycoprotein Folding Intermediates.

*J.* *Biol*. Chem. 279:46280–46285. doi:10.1074/JBC.M408404200.

Carvalho, P., V. Goder, and T.A. Rapoport. 2006. Distinct ubiquitin-ligase complexes define convergent pathways for the degradation of ER proteins. Cell. 126:361–373.

Chiapparino, A., A. Grbavac, H.R.A. Jonker, Y. Hackmann, S. Mortensen, E. Zatorska, A. Schott, G. Stier, K. Saxena, K. Wild, H. Schwalbe, S. Strahl, and I. Sinning. 2020. Functional implications of MIR domains in protein O-mannosylation. Elife. 9:1–23. doi:10.7554/ELIFE.61189.

Clerc, S., C. Hirsch, D.M. Oggier, P. Deprez, C. Jakob, T. Sommer, and M. Aebi. 2009. Htm1 protein generates the N-glycan signal for glycoprotein degradation in the endoplasmic reticulum. J Cell Biol. 184:159–172.

Denic, V., E.M. Quan, and J.S. Weissman. 2006. A luminal surveillance complex that selects misfolded glycoproteins for ER-associated degradation. Cell. 126:349–359.

Ellgaard, L., and A. Helenius. 2003. Quality control in the endoplasmic reticulum. Nat Rev Mol Cell Biol. 4:181–191.

Fujita, M., O.T. Yoko, and Y. Jigami. 2006. Inositol deacylation by Bst1p is required for the quality control of glycosylphosphatidylinositol-anchored proteins. Mol Biol Cell. 17:834–850.

Fukuda, S., M. Sumii, Y. Masuda, M. Takahashi, N. Koike, J. Teishima, H. Yasumoto, T. Itamoto, T. Asahara, K. Dohi, and K. Kamiya. 2001. Murine and human SDF2L1 is an endoplasmic reticulum stress-inducible gene and encodes a new member of the Pmt/rt protein family. Biochem Biophys Res Commun. 280:407–414.

Goder, V., and A. Melero. 2011. Protein O-mannosyltransferases participate in ER protein quality control. J Cell Sci. 124:144–153.

de Groot, P.W.J., K.J. Hellingwerf, and F.M. Klis. 2003. Genome-wide identification of fungal GPI proteins. Yeast. 20:781–796. doi:10.1002/YEA.1007.

Hammond, C., I. Braakman, and A. Helenius. 1994. Role of N-linked oligosaccharide recognition, glucose trimming, and calnexin in glycoprotein folding and quality control. Proc Natl Acad Sci U S A. 91:913–917.

Harty, C., S. Strahl, and K. Romisch. 2001. O-mannosylation protects mutant alpha-factor precursor from endoplasmic reticulum-associated degradation. Mol Biol Cell. 12:1093–1101.

Haselbeck, A., and W. Tanner. 1983. O-glycosylation in Saccharomyces cerevisiae is initiated at the endoplasmic reticulum. FEBS Lett. 158:335–338.

Helenius, A., and M. Aebi. 2004. Roles of N-Linked Glycans in the Endoplasmic Reticulum. 10.1146/annurev.biochem.73.011303.073752. 73:1019–1049. doi:10.1146/ANNUREV.BIOCHEM.73.011303.073752.

Hiller, M.M., A. Finger, M. Schweiger, and D.H. Wolf. 1996. ER degradation of a misfolded luminal protein by the cytosolic ubiquitin-proteasome pathway. Science (80*-. ).* 273:1725– 1728.

Hirayama, H., M. Fujita, T. Yoko-o, and Y. Jigami. 2008. O-mannosylation is required for degradation of the endoplasmic reticulum-associated degradation substrate Gas1*p via the ubiquitin/proteasome pathway in Saccharomyces cerevisiae. J Biochem. 143:555–567.

Jarosch, E., C. Taxis, C. Volkwein, J. Bordallo, D. Finley, D.H. Wolf, and T. Sommer. 2002. Protein dislocation from the ER requires polyubiquitination and the AAA-ATPase Cdc48. Nat Cell Biol. 4:134–139.

Jonikas, M.C., S.R. Collins, V. Denic, E. Oh, E.M. Quan, V. Schmid, J. Weibezahn, B. Schwappach, P. Walter, J.S. Weissman, and M. Schuldiner. 2009. Comprehensive characterization of genes required for protein folding in the endoplasmic reticulum. Science (80-. ). 323:1693–1697.

Kim, I., K. Mi, and H. Rao. 2004. Multiple interactions of rad23 suggest a mechanism for ubiquitylated substrate delivery important in proteolysis. Mol Biol Cell. 15:3357–3365.

Kostova, Z., and D.H. Wolf. 2005. Importance of carbohydrate positioning in the recognition of mutated CPY for ER-associated degradation. J Cell Sci. 118:1485–1492.

Krshnan, L., M.L. van de Weijer, and P. Carvalho. 2022. Endoplasmic Reticulum-Associated Protein Degradation. Cold Spring Harb. Perspect. Biol. 14. doi:10.1101/CSHPERSPECT.A041247.

Kushnirov, V. V. 2000. Rapid and reliable protein extraction from yeast. Yeast. 16:857– 860.

Larsen, I.S.B., Y. Narimatsu, H.J. Joshi, L. Siukstaite, O.J. Harrison, J. Brasch, K.M. Goodman, L. Hansen, L. Shapiro, B. Honig, S.Y. Vakhrushev, H. Clausen, and A. Halim. 2017. Discovery of an O-mannosylation pathway selectively serving cadherins and protocadherins. Proc. Natl. Acad. Sci. U. S. A. 114:11163–11168. doi:10.1073/PNAS.1708319114.

Lemoine, F., D. Correia, V. Lefort, O. Doppelt-Azeroual, F. Mareuil, S. Cohen-Boulakia, and O. Gascuel. 2019. NGPhylogeny.fr: new generation phylogenetic services for non-specialists. Nucleic Acids Res. 47:W260–W265. doi:10.1093/NAR/GKZ303.

Leto, D.E., D.W. Morgens, L. Zhang, C.P. Walczak, J.E. Elias, M.C. Bassik, and R.R. Kopito. 2019. Genome-wide CRISPR Analysis Identifies Substrate-Specific Conjugation Modules in ER-Associated Degradation. Mol. Cell. 73:377–389.e11. doi:10.1016/J.MOLCEL.2018.11.015.

Liu, J., and A. Mushegian. 2003. Three monophyletic superfamilies account for the majority of the known glycosyltransferases. Protein Sci. 12:1418–1431. doi:10.1110/PS.0302103.

Lommel, M., and S. Strahl. 2009. Protein O-mannosylation: conserved from bacteria to humans. Glycobiology. 19:816–828.

Meyer, H.J., and M. Rape. 2014. Enhanced protein degradation by branched ubiquitin chains. Cell. 157:910–921.

Molinari, M., K.K. Eriksson, V. Calanca, C. Galli, P. Cresswell, M. Michalak, and A. Helenius. 2004. Contrasting Functions of Calreticulin and Calnexin in Glycoprotein Folding and ER Quality Control. Mol. Cell. 13:125–135. doi:10.1016/S1097-2765(03)00494-5.

Naik, R.R., and E.W. Jones. 1998. The PBN1 gene of Saccharomyces cerevisiae: an essential gene that is required for the post-translational processing of the protease B precursor. Genetics. 149:1277–1292. doi:10.1093/GENETICS/149.3.1277.

Nakatsukasa, K., G. Huyer, S. Michaelis, and J.L. Brodsky. 2008. Dissecting the ER-associated degradation of a misfolded polytopic membrane protein. Cell. 132:101–112.

Nakatsukasa, K., S. Okada, K. Umebayashi, R. Fukuda, S. Nishikawa, and T. Endo. 2004. Roles of O-Mannosylation of Aberrant Proteins in Reduction of the Load for Endoplasmic Reticulum Chaperones in Yeast. J Biol Chem. 279:49762–49772.

Neubert, P., A. Halim, M. Zauser, A. Essig, H.J. Joshi, E. Zatorska, I.S.B. Larsen, M. Loibl, J. Castells-Ballester, M. Aebi, H. Clausen, and S. Strahl. 2016. Mapping the O-Mannose glycoproteome in saccharomyces cerevisiae. Mol. Cell. Proteomics. 15:1323–1337. doi:10.1074/MCP.M115.057505/.

Nuoffer, C., A. Horvath, and H. Riezman. 1993. Analysis of the sequence requirements for glycosylphosphatidylinositol anchoring of Saccharomyces cerevisiae Gas1 protein. J Biol Chem. 268:10558–10563.

Quan, E.M., Y. Kamiya, D. Kamiya, V. Denic, J. Weibezahn, K. Kato, and J.S. Weissman. 2008. Defining the glycan destruction signal for endoplasmic reticulum-associated degradation. Mol Cell. 32:870–877.

Ribeiro, C.V., B.F.B. Rocha, D.D.S. Moreira, V. Peruhype-Magalhães, and S.M.F. Murta. 2019. Mannosyltransferase (GPI-14) overexpression protects promastigote and amastigote forms of Leishmania braziliensis against trivalent antimony. Parasites and Vectors. 12:1–7. doi:10.1186/S13071-019-3305-2/TABLES/2.

Ruiz-Canada, C., D.J. Kelleher, and R. Gilmore. 2009. Cotranslational and Posttranslational N-Glycosylation of Polypeptides by Distinct Mammalian OST Isoforms. Cell. 136:272–283. doi:10.1016/J.CELL.2008.11.047.

Satpute-Krishnan, P., M. Ajinkya, S. Bhat, E. Itakura, R.S. Hegde, and J. Lippincott-Schwartz. 2014. ER stress-induced clearance of misfolded GPI-anchored proteins via the secretory pathway. Cell. 158:522–533.

Sikorska, N., L. Lemus, A. Aguilera-Romero, J. Manzano-Lopez, H. Riezman, M. Muniz, and V. Goder. 2016. Limited ER quality control for GPI-anchored proteins. J Cell Biol. 213:693– 704.

Spear, E.D., and D.T. Ng. 2005. Single, context-specific glycans can target misfolded glycoproteins for ER-associated degradation. J Cell Biol. 169:73–82.

Stein, A., A. Ruggiano, P. Carvalho, and T.A. Rapoport. 2014. Key Steps in ERAD of Luminal ER Proteins Reconstituted with Purified Components. Cell. 158:1375–1388.

Subramanian, S., C.A. Woolford, E. Drill, M. Lu, and E.W. Jones. 2006. Pbn1p: An essential endoplasmic reticulum membrane protein required for protein processing in the endoplasmic reticulum of budding yeast. Proc. Natl. Acad. Sci. U. S. A. 103:939–944. doi:10.1073/pnas.0505570103.

Swanson, R., M. Locher, and M. Hochstrasser. 2001. A conserved ubiquitin ligase of the nuclear envelope/endoplasmic reticulum that functions in both ER-associated and Matalpha2 repressor degradation. Genes Dev. 15:2660–2674.

Travers, K.J., C.K. Patil, L. Wodicka, D.J. Lockhart, J.S. Weissman, and P. Walter. 2000. Functional and genomic analyses reveal an essential coordination between the unfolded protein response and ER-associated degradation. Cell. 101:249–258.

Wang, Y., Y. Maeda, Y.S. Liu, Y. Takada, A. Ninomiya, T. Hirata, M. Fujita, Y. Murakami, and T. Kinoshita. 2020. Cross-talks of glycosylphosphatidylinositol biosynthesis with glycosphingolipid biosynthesis and ER-associated degradation. Nat. Commun. 11. doi:10.1038/S41467-020-14678-2.

Weill, U., I. Yofe, E. Sass, B. Stynen, D. Davidi, J. Natarajan, R. Ben-Menachem, Z. Avihou, O. Goldman, N. Harpaz, S. Chuartzman, K. Kniazev, B. Knoblach, J. Laborenz, F. Boos, J. Kowarzyk, S. Ben-Dor, E. Zalckvar, J.M. Herrmann, R.A. Rachubinski, O. Pines, D. Rapaport, S.W. Michnick, E.D. Levy, and M. Schuldiner. 2018. Genome-wide SWAp-Tag yeast libraries for proteome exploration. Nat. Methods. 15:617–622. doi:10.1038/s41592-018-0044-9.

Werner, E.D., J.L. Brodsky, and A.A. McCracken. 1996. Proteasome-dependent endoplasmic reticulum-associated protein degradation: an unconventional route to a familiar fate. Proc Natl Acad Sci U S A. 93:13797–13801.

Wiggins, C.A.R., and S. Munro. 1998. Activity of the yeast MNN1 α-1,3- mannosyltransferase requires a motif conserved in many other families of glycosyltransferases. Proc. Natl. Acad. Sci. U. S. A. 95:7945–7950.

Xiang, M.-H., X.-X. Xu, C.-D. Wang, S. Chen, S. Xu, X.-Y. Xu, N. Dean, N. Wang, and X.-D. Gao. 2022. Topological and enzymatic analysis of human Alg2 mannosyltransferase reveals its role in lipid-linked oligosaccharide biosynthetic pathway. doi:10.1038/s42003-022-03066-9.

Xu, C., S. Wang, G. Thibault, and D.T.W. Ng. 2013. Futile Protein Folding Cycles in the ER Are Terminated by the Unfolded Protein O-Mannosylation Pathway. Science (80*-. ).* 340:978 LP-- 981.

