## Supplementary material for "ER-associated degradation relying on protein O-mannosylation": suppl figures Lemus et al

Supplementary Figures S1-S5 and legends

suppl. Figure S1

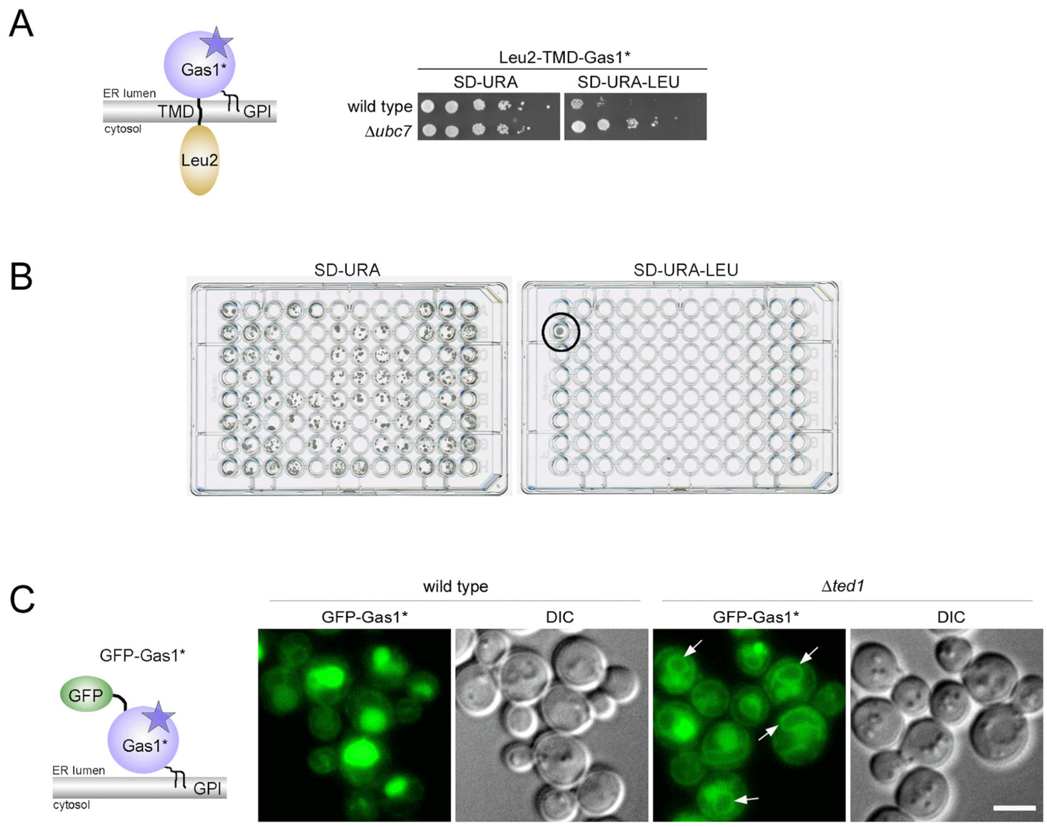


**Experimental data linked to Figure 1. (A) Design and functionality test of the first screen.** Wild type cells and cells with *UBC7* deleted *(Δubc7*) were used to express the plasmid-borne reporter construct Leu2-TMD-Gas1* (graphical depiction). The plasmid contained *URA3* as a marker and expressed the reporter from the *GAL4* promoter. Cells were grown in medium lacking uracil and spotted in serial dilutions in equal concentration on plates containing synthetic medium lacking uracil (SD-URA) or lacking both uracil and leucine (SD-URA-LEU). Plates were incubated for three days (SD-URA) or five days (SD-URA-LEU) at 30°C prior to imaging. The growth of *Δubc7* mutant cells in the absence of leucine (SD-URA-LEU) indicated an impairment in reporter degradation, as expected. **(B) Screen readout.** Representative example for pairs of 96-well plates used for the identification of hits. Transformants of the yeast deletion collection (*Δxxx*) and the collection of decreased abundance by mRNA perturbation (*DAmP*) alleles were first grown in 96-well plates containing liquid synthetic medium lacking uracil (SD-URA). Note that the organization of the libraries is such that not all wells contain cells. After growth in SD-URA, cells were replica-plated into 96-well plates containing medium lacking uracil (SD-URA) and lacking both uracil and leucine (SD-URA-LEU), a pair of plates is shown as example. Growth in both media was considered a hit (circled well). **(C) Design of the second screen.** Wild-type cells and cells with *TED1* deleted *(Δted1*) were used to express genomic N-terminally GFP-tagged Gas1* (graphical depiction) for live cell high-throughput fluorescence microscopy and differential interference contrast (DIC) microscopy. Ted1 is involved in GPI anchor remodeling. Its deletion was known to reduce the ER export and vacuolar targeting of Gas1* and to increase its routing to ERAD (Sikorska et al., 2016). In agreement, while most GFP was visible inside vacuoles in wild-type cells, indicating efficient targeting of GFP-Gas1* to vacuoles, a significant fraction of the protein accumulated in the perinuclear ER in Δ*ted1* cells (arrows). Differential interference contrast (DIC) microscopy was used to identify vacuoles. Scale bar: 4µm.

suppl. Figure S2

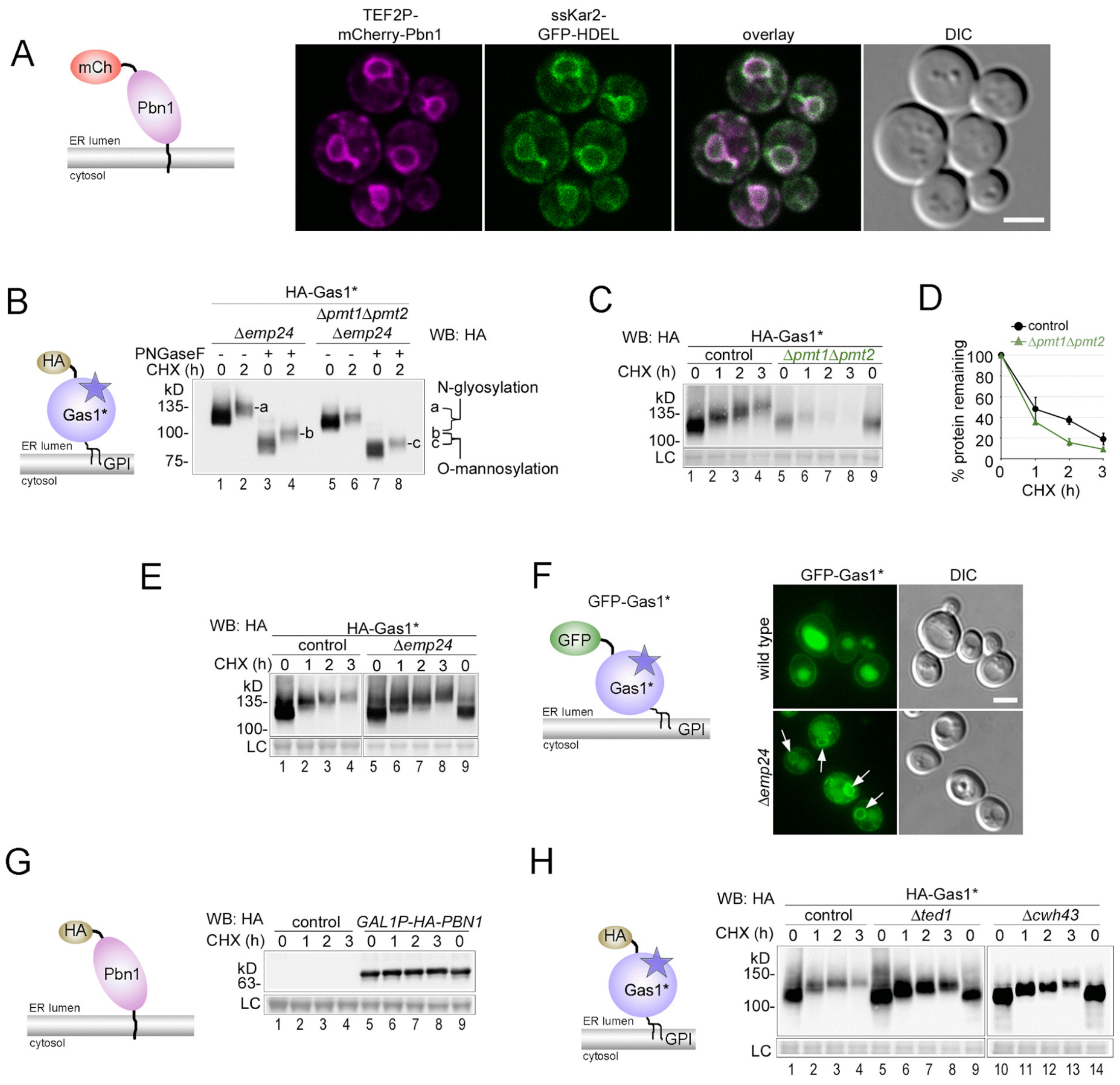


**Experimental data linked to Figure 2. (A) Pbn1 localizes to the ER.** Yeast cells co-expressing genomic N-terminally mCherry-tagged Pbn1 from the strong constitutive *TEF2* promoter (graphical depiction) and the ER marker ssKar2-GFP-HDEL were analyzed by live cell fluorescence microscopy and DIC microscopy. The degree of colocalization indicated that Pbn1 does not "leak" from the ER, even at a high expression level. Scale bar: 3μm. **(B) Gas1* is N-glycosylated and O-mannosylated.** Cells with the indicated deletions were used to express plasmid-borne HA-tagged Gas1* (graphical depiction) for cycloheximide (CHX) shut-off experiments. To prevent rapid ER export and vacuolar degradation of Gas1* upon deletion of *PMT1* and *PMT2* (*ΔpmtΔpmt2*), the experiment was performed in the *Δemp24* background (suppl. Fig. 2C, and (Sikorska et al., 2016)). Cells were lysed at indicated time points and, where indicated, lysates were treated with Peptide-N-Glycosidase F (PNGaseF) for the removal of all N-linked glycans, prior to SDS-PAGE and Western blot (WB) analysis with anti-HA antibodies. The visible shift in MW after treatment with PNGaseF was indicative of extensive Gas1* N-glycosylation (compare lanes 2 and 4). The additional shift in MW visible in *Δpmt1Δpmt2Δemp24* cells compared to *Δemp24* cells was indicative of protein O-mannosylation (compare lanes 4 and 8). **(C)-(D) Gas1* degradation is accelerated in the absence of the Pmt1/2-complex.** Control cells and *Δpmt1Δpmt2* cells were used to express plasmid-borne HA-tagged Gas1* for CHX shut-off experiments. Cells were lysed at indicated time points after addition of CHX and remaining HA-Gas1* was measured by SDS-PAGE and WB with anti-HA antibodies. Membrane staining with Ponceau served as loading control (LC). Cells with *PMT1* and *PMT2* deleted (*Δpmt1Δpmt2*) showed two effects. First, the increase in MW was drastically reduced (compare lanes 1-4 with 5-9), in agreement with the protein being O-mannosylated primarily by the Pmt1/2-complex. Second, Gas1* was degraded faster (compare lanes 1-4 with 5-9, and graph). Faster degradation was due to a loss of ER retention and increased routing to the vacuole under these conditions, in agreement with the Pmt1/2-complex mediating protein O-mannosylation of Gas1* in combination with ER retention (Goder and Melero, 2011; Sikorska et al., 2016). **(E) Gas1* O-mannosylation is not reduced or eliminated in *Δemp24* cells.** Control cells and *Δemp24* cells were used to express plasmid-borne HA-tagged Gas1* for CHX shut-off experiments. Cells with the ER export factor *EMP24* deleted showed comparable increase in MW of Gas1* during the chase period. Stabilization of Gas1*in *Δemp24* cells was reported previously (Sikorska et al., 2016). **(F) Gas1* O-mannosylation occurs inside the ER.** Wild-type cells and *Δemp24* cells were used to express genomic N-terminally GFP-tagged Gas1* (graphical depiction) for live cell fluorescence microscopy and DIC microscopy. Arrows indicate accumulated GFP-Gas1* inside the ER in *Δemp24* cells. Since the increase in molecular weight (MW) is comparable to that in control cells (suppl. Fig. S2E), it indicates that protein O-mannosylation occurs within the ER. Scale bar: 3µm. **(G) Stability of HA-tagged Pbn1 expressed from the *GAL1* promoter**. Control cells and cells expressing genomic N-terminally HA-tagged Pbn1 from the *GAL1* promoter (graphical depiction) were used for CHX shut-off experiments. Cells were grown in medium containing galactose as carbon source. Overexpressed Pbn1 remained stable throughout the experimental period. This strain background was used for all experiments using overexpression of HA-Pbn1. **(H) Gas1* O-mannosylation remains unaffected in mutants that impact GPI metabolism.** Control cells and cells with the indicated deletions were used to express plasmid-borne HA-tagged Gas1* for CHX shut-off experiments. The deletion mutants showed an increase in MW of Gas1* during the chase period comparable to that of control cells. Stabilization of Gas1*in these deletion mutants was reported previously (Sikorska et al., 2016).

suppl. Figure S3

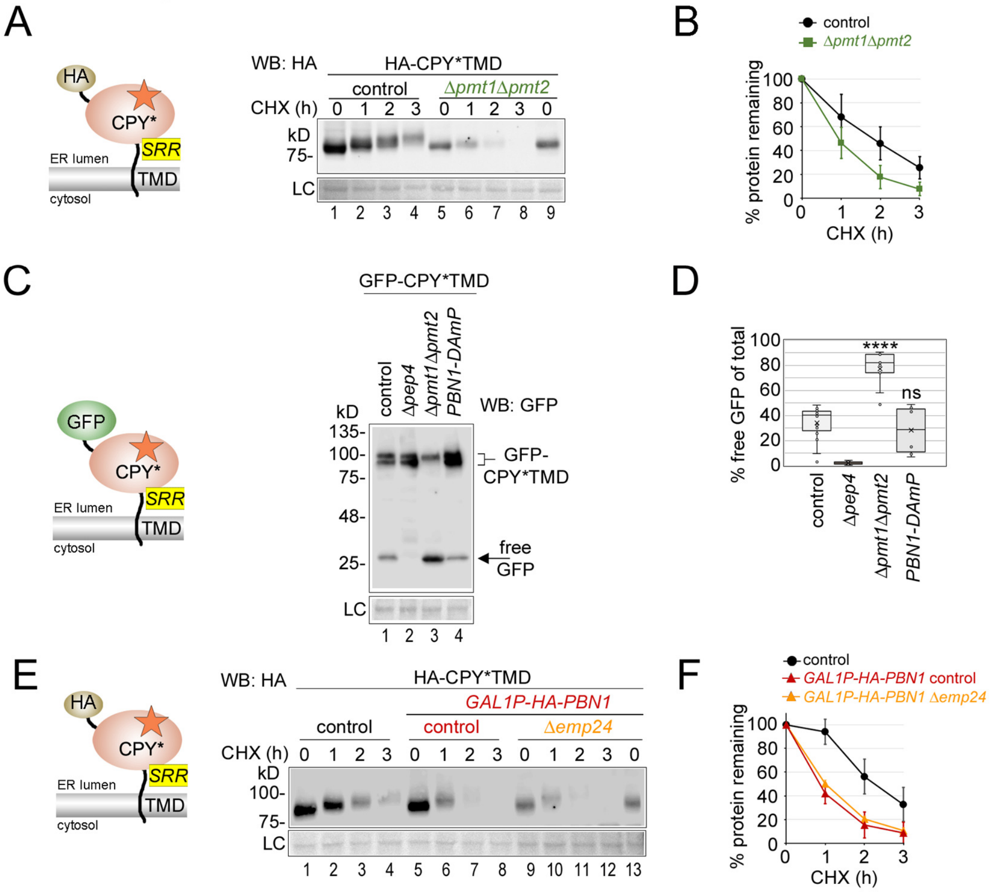


**Experimental data linked to Figure 3. (A)-(B) CPY*TMD is O-mannosylated.** Control cells and cells with *PMT1* and *PMT2* deleted (*ΔpmtΔpmt2*) were used to express plasmid-borne HA-tagged CPY*TMD (graphically depicted) for CHX shut-off experiments. Cells were lysed at indicated time points after addition of CHX and remaining protein was measured by SDS-PAGE and WB with anti-HA antibodies. Like seen with Gas1*, CPY*TMD showed two phenotypes when expressed in *ΔpmtΔpmt2* cells. First, the increase in MW was drastically reduced (compare lanes 1-4 with 5-9), consistent with the protein being O-mannosylated by the Pmt1/2-complex. Second, CPY*TMD was degraded faster (compare lanes 1-4 with 5-9, and graph). Faster degradation was due to a loss of ER retention and increased routing to the vacuole under these conditions (see C-D). **(C)-(D) Faster degradation of CPY*TMD in *Δpmt1Δpmt2* cells correlates with increased routing to the vacuole.** Faster degradation of CPY*TMD in *Δpmt1Δpmt2* cells resembled results obtained with Gas1* (suppl. Figs. S2C and D) and was tested for increased targeting to the vacuole in this background. To perform a comparative analysis for the amount of CPY*TMD routed to the vacuole in different mutant backgrounds, we made use of the well-known GFP-cleavage assay (Klionsky et al., 2021). A GFP-tagged version of CPY*TMD was generated (graphical depiction), which led to the appearance of (free) GFP in WB analysis after vacuolar degradation of the protein due to the proteolytic stability of the GPF tag (lane 1, "free GFP"). Free GFP was generated inside the vacuole because it was not generated in the absence of the vacuolar master protease Pep4 (lane 2). The graph displays the percentage of free GFP relative to the total GFP signal in each lysate. In *Δpmt1Δpmt2* cells, the relative amount of free GFP was increased, indicating an elevated trafficking of GFP-CPY*TMD to the vacuole (lane 3 and graph). Statistical significance ****$p<0.00005$ (unpaired two-tailed Student's t-test). ns = not significant. Together these findings agree with previous reports that the Pmt1/2-complex not only affects protein O-mannosylation but also ER retention, either directly or indirectly (Goder and Melero, 2011). Loss of ER retention of misfolded proteins can lead to an increase in efficient vacuolar degradation. **(E)-(F) CPY*TMD degradation kinetics upon Pbn1 overexpression was unaffected in *Δemp24* cells.** Control cells lacking Pbn1 overexpression, along with control and *Δemp24* cells expressing HA-tagged genomic Pbn1 from the *GAL1* promoter (*GAL1P-PBN1*), were used to express plasmid-borne HA-tagged CPY*TMD (as depicted) for CHX shut-off experiments in medium with galactose as carbon source. Cells were lysed at indicated time points after addition of CHX and remaining HA-CPY*TMD was measured by SDS-PAGE and WB with anti-HA antibodies. In contrast to the results obtained with deleted ERAD components, deletion of *EMP24* did not affect degradation of HA-CPY*TMD (lanes 5-12).

suppl. Figure S4

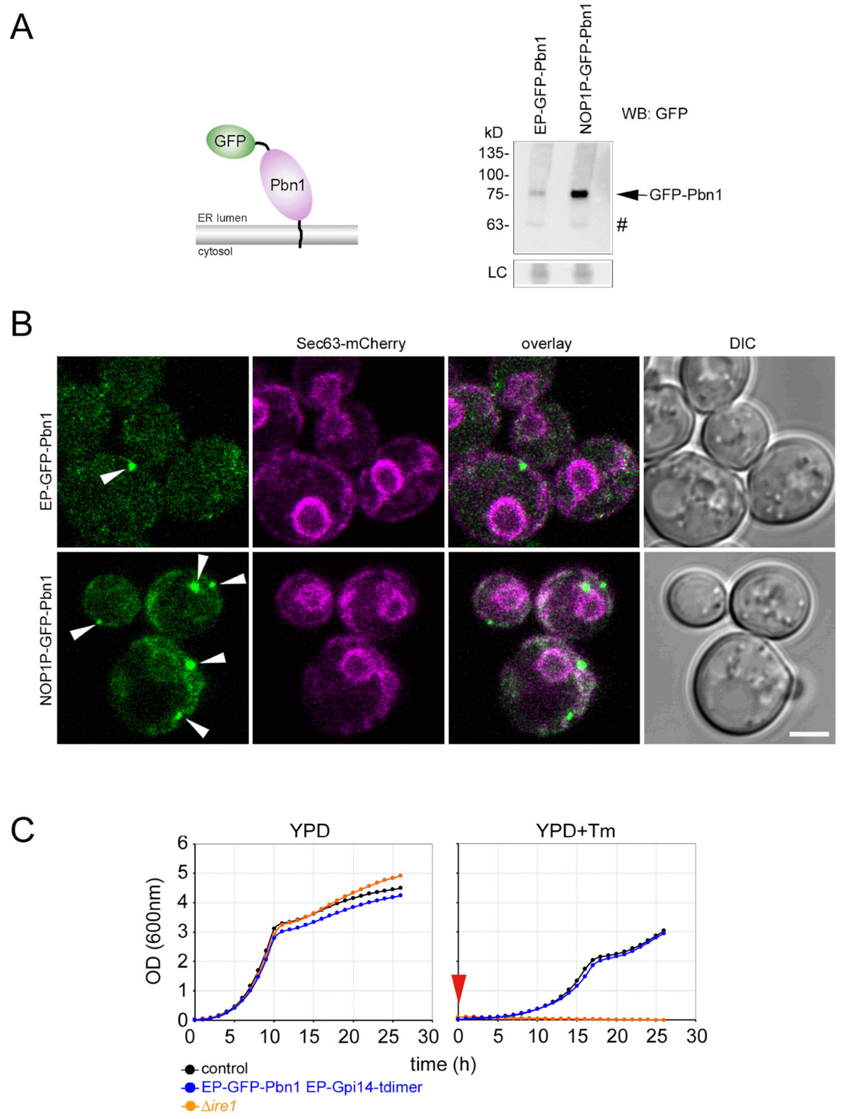


**Experimental data linked to Figure 4. (A) Comparison of cellular levels of GFP-tagged Pbn1 expressed from different promoters.** Expression of genomic N-terminally GFP-tagged Pbn1 (graphical depiction) from either the endogenous promoter (EP) or from the moderate *NOP1* promoter (NOP1P). Equal amounts of cells were taken for lysis and expression levels were compared by SDS-PAGE followed by WB analysis with antibodies against GFP. Hashtag indicates an unspecific band. The results indicated weak expression of GFP-Pbn1 from its endogenous promoter. **(B) Cellular localization of GFP-Pbn1.** Cell co-expressing genomic GFP-Pbn1 from different promoters like in (A) and the ER marker Sec63-mCherry were analyzed by live cell confocal fluorescence microscopy in combination with DIC microscopy. Note that N-terminal tagging of Pbn1 with GFP led to the concentration of the protein in cellular puncta (arrowheads) regardless of protein expression levels. Co-localization with the ER marker Sec63 suggested that Pbn1 punta localized to the ER. Scale bar: 2µm. **(C) Functionality tests of tagged versions of Pbn1 and Gpi14.** Wild type cells (control) and strains expressing genome-integrated fluorescently tagged Pbn1 and Gpi14, were grown in complete synthetic media (SD) in absence or presence of 1µg/ml tunicamycin (Tm) at 30°C. The drug was applied immediately after starting the experiment by resetting cellular density (red arrowhead). Growth was determined by automated measuring of absorbance of the individual cell cultures at 600nm over a period of 25 hours. Cells with *IRE1* deleted (*Δire1*) are known to be sensitive to the induction of ER stress with tunicamycin and were used as control. The strains co-expressing GFP-Pbn1 and Gpi14-tdimer showed no growth defect in SD medium in absence or presence of tunicamycin compared to the control strain. Since both proteins, Pbn1 and Gpi14, are essential genes, these results indicated that the utilized tagging did not compromise the essential functions of either protein.

suppl. Figure S5

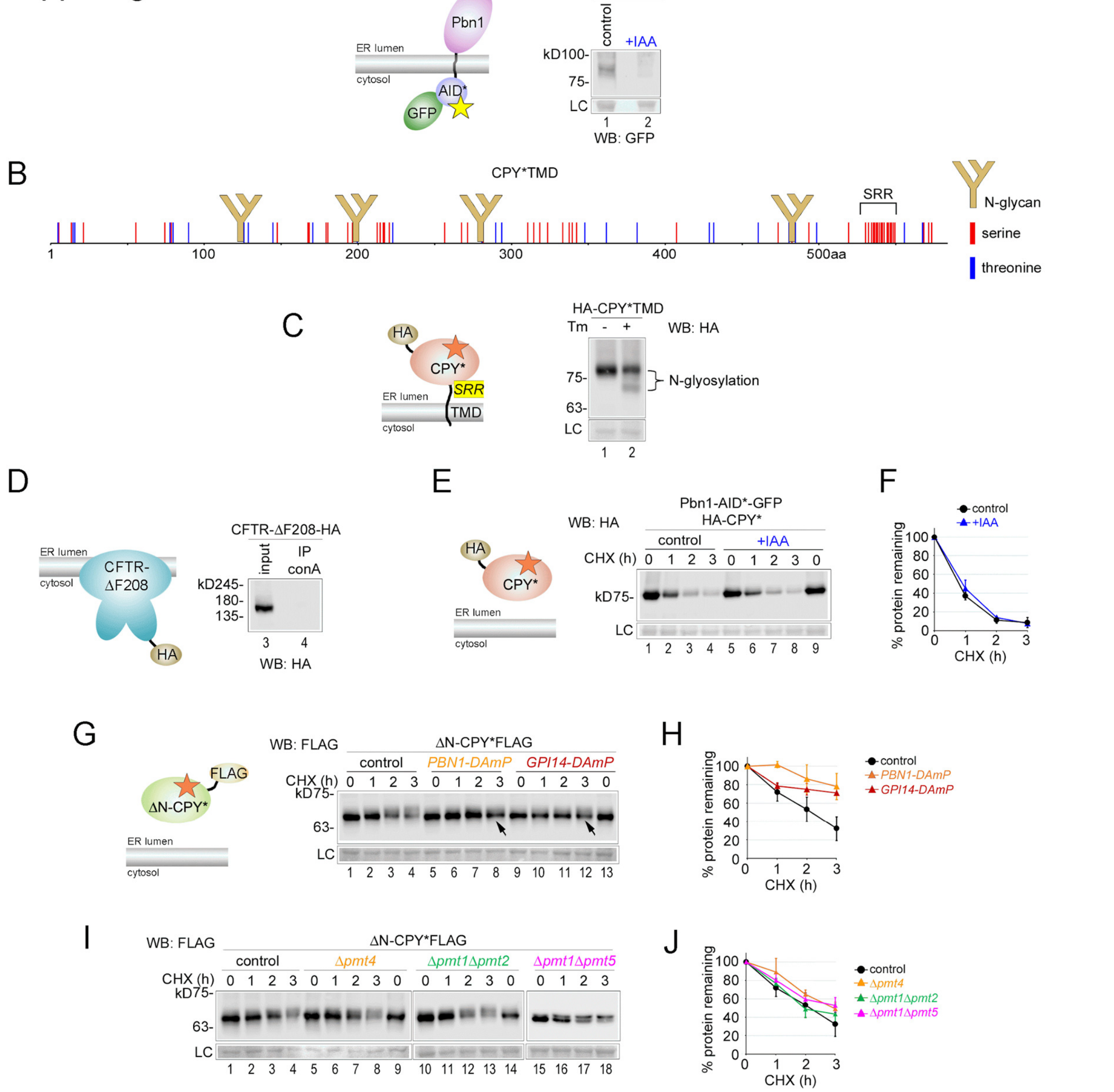


**Experimental data linked to Figure 5. (A) Auxin-induced degradation of Pbn1-AID*GFP.** Cells expressing genomic Pbn1 C-terminally tagged with the degron AID* in combination with GFP (graphical depiction) were supplemented with auxin (indole-3-acetic acid (IAA)) or only with solvent (control). One hour after addition of auxin cells were lysed followed by SDS-PAGE and WB analysis with antibodies against GFP. Pbn1-AID*GFP was no longer detectable by WB after induced degradation. **(B) A schematic of the protein sequence of CPY*TMD.** Like CPY*, CPY*TMD is modified with four N-glycans at the indicated positions. The C-terminal serine-rich region provides a unique motif that differs from the scattered serines and threonines found throughout the rest of the sequence and provides an explanation for why CPY*TMD, unlike CPY*, was O-mannosylated even in the presence of N-glycans. **(C) CPY*TMD is N-glycosylated.** Lysate from cells expressing HA-tagged CPY*TMD was analyzed by WB analysis before and after cells were grown in presence of 1µg/ml tunicamycin (Tm) for one hour. Newly synthesized CPY*TMD in presence of tunicamycin displayed a lower MW, demonstrating that the protein is normally N-glycosylated. **(D) Control for concanavalin-A-mediated pulldown.** Cells expressing plasmid-borne CFTR-ΔF208-HA (graphical depiction) were lysed, and the lysate was incubated with concanavalin-A-coupled beads (IP conA) for three hours, followed by SDS-PAGE and WB analysis with antibodies against HA. CFTR-ΔF208-HA is not glycosylated in yeast and its failure to be precipitated with conA served as a specificity control for the assay and validated the results obtained with ΔN-CPY* (Fig. 5H). **(E)-(F) Pbn1 does not affect the degradation of CPY*.** Cells expressing genomic Pbn1 C-terminally tagged with the degron AID* in combination with GFP were used to express plasmid-borne CPY*HA (graphical depiction) for CHX shut-off experiments. Five hours prior to the addition of CHX, cells were supplemented with auxin (IAA) or only with solvent (control). Mean values and standard deviations from two individual experiments are shown in the graph. **(G)-(H) Pbn1 and Gpi14 affect O-mannosylation and degradation of ΔN-CPY*.** Control cells and cells containing the *PBN1-DAmP* or *GPI14-DAmP* alleles were used to express plasmid-borne ΔN-CPY*FLAG (graphical depiction) for CHX shut-off experiments. The arrows indicate the accumulation of the population of ΔN-CPY*FLAG with lower MW. Mean values and standard deviations from at least three individual experiments are shown in the graph. Total remaining protein (upper and lower bands combined) has been quantified. The results revealed a stabilization of ΔN-CPY*FLAG in *PBN1-DAmP* and *GPI14-DAmP* cells, in agreement with the results obtained after auxin-induced degradation of Pbn1 or Gpi14 (Fig 5 I-L). **(I)-(J) Single and double deletion mutants of canonical PMTs did not affect O-mannosylation or degradation of ΔN-CPY*.** Yeast expresses six verified PMTs, with the most well-characterized being the conserved Pmt1, Pmt2 and Pmt4 (Lommel and Strahl, 2009). Control cells and cells with the indicated single and double deletions of PMT genes were used to express plasmid-borne ΔN-CPY*FLAG for CHX shut-off experiments. Neither in the *Δpmt4* mutant, which lacks the homodimeric Pmt4 complex, nor in the *Δpmt1Δpmt2* double mutant, which lacks the heterodimeric Pmt1/2 complex, did we detect alterations in turnover or relative O-mannosylation of ΔN-CPY*, i.e., no accumulation of the band with lower MW (lanes 1-14, and J). The same results were obtained with the *Δpmt1Δpmt5* mutant which simultaneously prevented the formation of the canonical and a cross-combinatorial Pmt1/2 complex (lanes 15-18, and J). Total remaining protein (upper and lower bands combined) has been quantified Mean values and standard deviations from at least two individual experiments are shown in the graph.

**References to supplemental data**


Goder, V., and A. Melero. 2011. Protein O-mannosyltransferases participate in ER protein quality control. *J Cell Sci*. 124:144–153.

Klionsky, D.J., A.K. Abdel-Aziz, S. Abdelfatah, M. Abdellatif, A. Abdoli, S. Abel, H. Abeliovich, M.H. Abildgaard, Y.P. Abudu, A. Acevedo-Arozena, I.E. Adamopoulos, K. Adeli, T.E. Adolph, A. Adornetto, E. Aflaki, G. Agam, A. Agarwal, B.B. Aggarwal, M. Agnello, P. Agostinis, J.N. Agrewala, A. Agrotis, P. V. Aguilar, S.T. Ahmad, Z.M. Ahmed, U. Ahumada-Castro, S. Aits, S. Aizawa, Y. Akkoc, T. Akoumianaki, H.A. Akpinar, A.M. Al-Abd, L. Al-Akra, A. Al-Gharaibeh, M.A. Alaoui-Jamali, S. Alberti, E. Alcocer-Gómez, C. Alessandri, M. Ali, M.A. Alim Al-Bari, S. Aliwaini, J. Alizadeh, E. Almacellas, A. Almasan, A. Alonso, G.D. Alonso, N. Altan-Bonnet, D.C. Altieri, É.M.C. Álvarez, S. Alves, C. Alves da Costa, M.M. Alzaharna, M. Amadio, C. Amantini, C. Amaral, S. Ambrosio, A.O. Amer, V. Ammanathan, Z. An, S.U. Andersen, S.A. Andrabi, M. Andrade-Silva, A.M. Andres, S. Angelini, D. Ann, U.C. Anozie, M.Y. Ansari, P. Antas, A. Antebi, Z. Antón, T. Anwar, L. Apetoh, N. Apostolova, T. Araki, Y. Araki, K. Arasaki, W.L. Araújo, J. Araya, C. Arden, M.A. Arévalo, S. Arguelles, E. Arias, J. Arikkath, H. Arimoto, A.R. Ariosa, D. Armstrong-James, L. Arnauné-Pelloquin, A. Aroca, D.S. Arroyo, I. Arsov, R. Artero, D.M.L. Asaro, M. Aschner, M. Ashrafizadeh, O. Ashur-Fabian, A.G. Atanasov, A.K. Au, P. Auberger, et al. 2021. Guidelines for the use and interpretation of assays for monitoring autophagy (4th edition)1. *Autophagy*. 17:1–382. doi:10.1080/15548627.2020.1797280.

Lemoine, F., D. Correia, V. Lefort, O. Doppelt-Azeroual, F. Mareuil, S. Cohen-Boulakia, and O. Gascuel. 2019. NGPhylogeny.fr: new generation phylogenetic services for non-specialists. *Nucleic Acids Res.* 47:W260–W265. doi:10.1093/NAR/GKZ303.

Sikorska, N., L. Lemus, A. Aguilera-Romero, J. Manzano-Lopez, H. Riezman, M. Muniz, and V. Goder. 2016. Limited ER quality control for GPI-anchored proteins. *J Cell Biol*. 213:693–704.
